# ModiFinder: Tandem Mass Spectral Alignment Enables Structural Modification Site Localization

**DOI:** 10.1101/2024.02.17.580849

**Authors:** Mohammad Reza Zare Shahneh, Michael Strobel, Giovanni Andrea Vitale, Christian Geibel, Yasin El Abiead, Neha Garg, Allegra T Aron, Vanessa V Phelan, Daniel Petras, Mingxun Wang

**Affiliations:** Department of Computer Science and Engineering, University of California Riverside, 900 University Ave., Riverside, CA, 92521, United States; Interfaculty Institute of Microbiology and Infection Medicine, University of Tuebingen, Auf der Morgenstelle 24, Tuebingen, 72076, Germany; Skaggs School of Pharmacy and Pharmaceutical Sciences, University of California San Diego, 9500 Gilman Dr., San Diego, CA, 92093, United States; School of Chemistry and Biochemistry, Georgia Institute of Technology, Atlanta,, 950 Atlantic Drive, Atlanta, GA, 30332, United States; Department of Chemistry and Biochemistry, University of Denver, 2101 East Wesley Ave, Denver, CO, 80210, United States; Department of Pharmaceutical Sciences, Skaggs School of Pharmacy and Pharmaceutical Sciences, University of Colorado, Anschutz Medical Campus, 12850 E Montview Blvd, Aurora, CO, 80045, United States; Department of Biochemistry, University of California Riverside, 900 University Ave., Riverside, CA, 92521, United States

**Author notes:** **Author Contributions**: MW conceptualized the idea. MW organized the project. MRZS implemented the software and performed the benchmarking. MRZS, MW, MS performed the analysis of the findings. MW and MRZS wrote the manuscript, MW and MRZS contributed to improvement ideas. GAV, CG and DP acquired LC-MS/MS data. YE contributed to improvement ideas and edited the manuscript. GAV, CG and DP inspected and manually interpreted MS/MS spectra. All authors edited and approved the final manuscript. Competing interest statement: MW is a co-founder of Ometa Labs.

## Abstract

Untargeted tandem mass spectrometry (MS/MS) has become a high-throughput method to measure small molecules in complex samples. One key goal is the transformation of these MS/MS spectra into chemical structures. Computational techniques such as MS/MS library search have enabled the re-identification of known compounds. Analog library search and molecular networking extend this identification to unknown compounds. While there have been advancements in metrics for the similarity of MS/MS spectra of structurally similar compounds, there is still a lack of automated methods to provide site specific information about structural modifications. Here we introduce ModiFinder that leverages the alignment of peaks in MS/MS spectra between structurally related known and unknown small molecules. Specifically, ModiFinder focuses on shifted MS/MS fragment peaks in the MS/MS alignment. These shifted peaks putatively represent substructures of the known molecule that contain the site of the modification. ModiFinder synthesizes these information together and scores the likelihood for each atom in the known molecule to be the modification site. We demonstrate in this manuscript how ModiFinder can effectively localize modifications which extends the capabilities of MS/MS analog searching and molecular networking to accelerate the discovery of novel compounds.

## Introduction

Tandem mass spectrometry (MS/MS) is a powerful analytical technique for identifying the structure of small molecules^1^. However, translating the MS/MS spectra to 2D chemical structures poses a significant challenge in the field^2^. Spectrum library matching^3^ is a key strategy within the field of metabolomics to annotate known compounds. However, in untargeted mass spectrometry experiments, on average 87% of MS/MS spectra remain unidentified by spectral library search^4^. To bridge this gap, modification aware spectral matching tools such as analog library search^5^ and molecular networking^6,7^ leverage the concept of structural propagation of known to unknown compounds - bridging between molecules with a conserved core structure but exhibit structural modifications^8–10^. A key shortcoming of these approaches is that they determine which pairs of MS/MS are putatively similar in structure, but do not describe explicitly the structural difference, leaving the manual interpretation up to chemists. To tackle this shortcoming, we have developed a computational approach, *ModiFinder,* that builds upon MS/MS matching and produces putative suggestions on the structural difference between known and unknown structural analogs.

Our approach borrows a concept from the computational challenge of site localization of post-translational modifications (PTM) of peptides in bottom-up proteomics^11–13^. In PTM site localization, b/y ions that f ank the modification site are used to localize the putative PTM on a linear peptide. Here, we translate this concept to localize structural modifications of small molecule graphs - representing 2D molecular structures. In contrast to PTM site localization, the ability to explain the MS/MS fragmentation, while simpler in peptides, is significantly more difficult in small molecules^14^. This complexity is underscored by the plethora of methodologies, including MetFrag^15^, MAGMa^16^, MIDAS^17^, and MS-Finder^18^ developed to tackle the small molecule fragmentation analysis. Despite the ongoing challenge of explaining MS/MS fragmentation^19^, we have found that these *in silico* approaches are still useful in addressing the problem of localizing structural modifications on small molecules.

ModiFinder leverages the insight that flanking masses for small molecule modifications can be determined by comparing the MS/MS spectrum of an unknown structure with a modification (MS2-unknown) with the MS/MS spectrum of the unmodified known structure (MS2-known). Specifically, the peaks that are shifted by the mass of the modification between the MS2-known and MS2-unknown putatively represent substructures that contain the modification site. Conversely, peaks that do not shift in mass between MS2-known and MS2-unknown, are less likely to include the modification (**Fig. 1**). Combining this information, ModiFinder computes a likelihood score for the specific site of modification across all atoms in the known compound (S-known). To accomplish this, each peak is assigned a set of possible substructures using combinatorial fragmentation^16^. Then, for each peak of MS2-known that has a corresponding shifted peak in MS2-unknown (signifying the substructure includes site of the modification), ModiFinder increases the likelihood scores of atoms in the assigned substructures for the shifted peaks. If the peak is unshifted, the likelihood is decreased. Finally, ModiFinder is able to map out the likelihood landscape for where the modification may occur across the S-known using the likelihood scores (**Fig. 2**).

**Figure 1:**
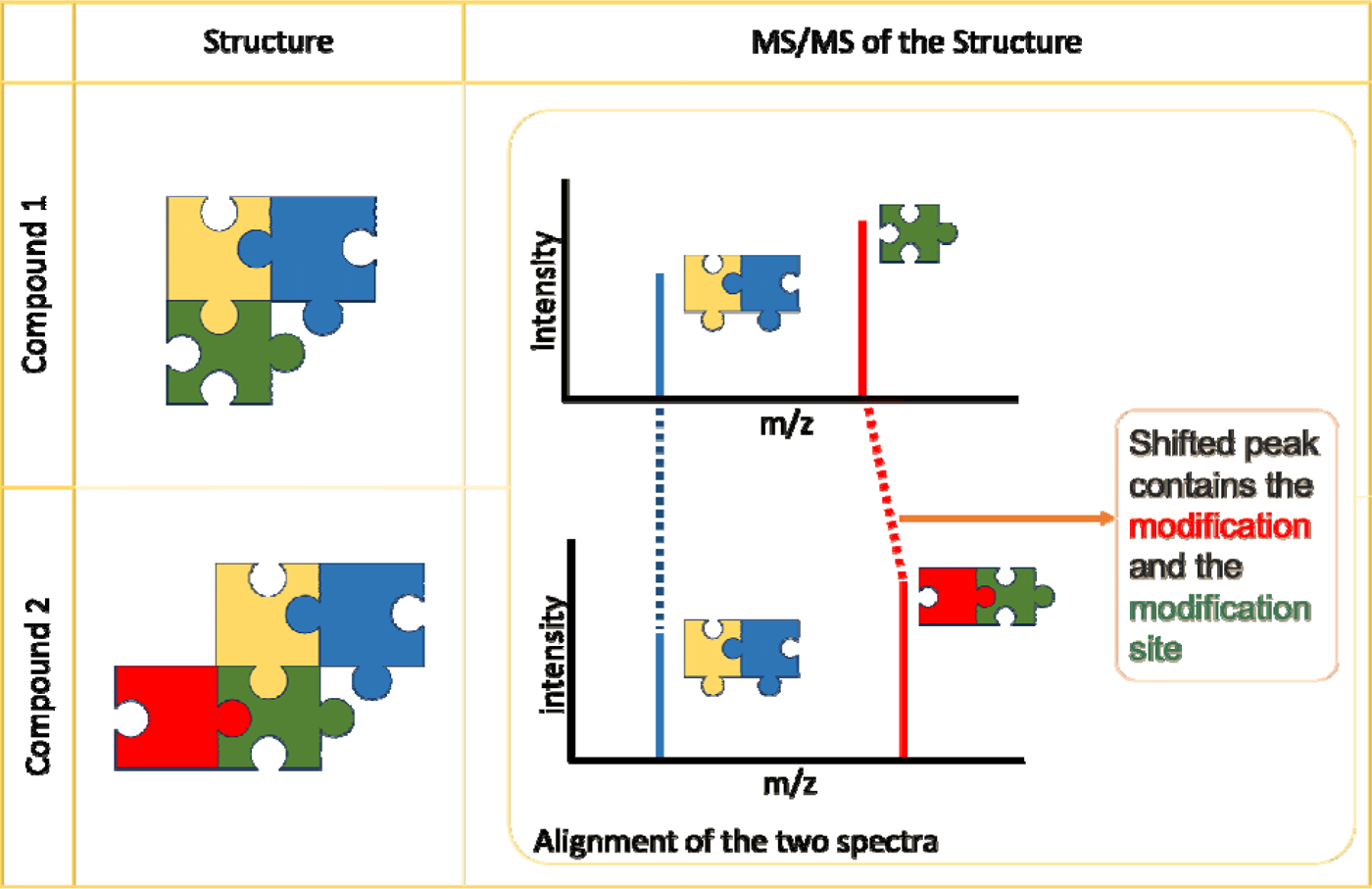
Illustration of the intuition behind ModiFinder. The MS/MS of Compound 1 and Compound 2 are aligned and matched peaks along with the substructures assigned to them are visualized. The matched peaks are shown in blue for the unshift peak and red for the shift peak. The matched shift peaks differ by the mass of the modification (the red puzzle piece) and contain the modification site (green puzzle piece).

**Figure 2:**
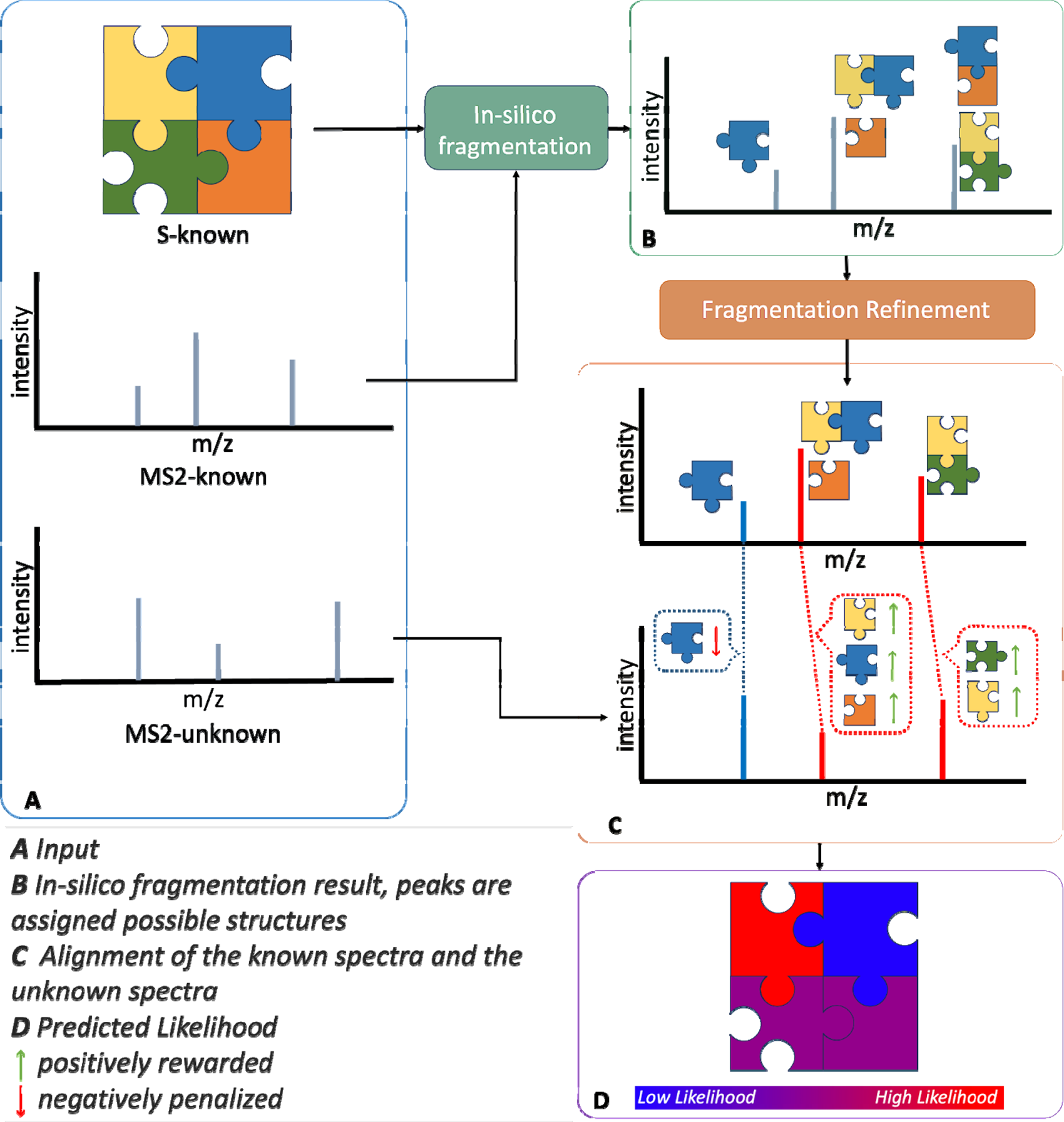
The Overview of ModiFinder algorithm. **A)** The input of ModiFinder, includes the known structure (S-known), spectra of the known compound (MS2-known), and spectra of the unknown compound (MS2-unknown). **B)** First, in-silico fragmentation methods compute potential substructure annotations for each MS/MS peak. Then, these substructures are refined by molecular formula and with helper MS/MS with a similar structure. **C)** Spectral alignment is performed to identify corresponding peaks (shifted and unshifted). The atoms in the substructures assigned to the shifted peaks are positively rewarded (increase in score) and atoms in the substructures assigned to the unshifted peaks are negatively penalized (decrease in score). **D)** Finally, a likelihood score is calculated proportionally to each atom’s score

We complement here the present of the ModiFinder approach with an evaluation of ModiFinder’s performance and limitations in identifying the modification site. Additionally, ModiFinder is presented as a command line tool and an interactive graphical web interface. Finally, we showcase empirical examples of how ModiFinder’s computational approach can be combined with domain knowledge expertise to facilitate the discovery of new natural products.

## Results and Discussion

### Benchmarking Data and Assessment Criteria

Pairs of structurally similar compounds with a single structural modification were used to assess the performance and accuracy of ModiFinder. These pairs were derived from the data available in four reference MS/MS libraries. In aggregate, the benchmark set contains 12,909 pairs with M+H adducts, that differ by a single structural modification, measured under the same experimental conditions, i.e., the same adduct and instrument (**See Data availability**). An additional filtering process was applied to these MS/MS pairs to only include pairs that have at least one shifted peak which can be explained by a substructure of the parent compound. After this filter, the majority (62% of the total pairs, 8033 pairs) of the pairs remain (**Fig. 7**).

Any evaluation metric that assesses the effectiveness of ModiFinder must strike a balance between two essential criteria: *proximity cover* and *ambiguity cover*.

**Proximity cover** assesses the distribution of likelihood scores relative to the true modification site and examines whether the high-scoring atoms are in close proximity to the actual modification site.

**Ambiguity cover** evaluates the entropy of the prediction array and its informativeness. For instance, an array where most atoms have the same high-score exhibits high ambiguity and may not be helpful for localization.

Several baseline metrics were considered but exhibited specific weaknesses. For example, if an evaluation function only checks if an algorithm assigns the highest score to the true modification site, an algorithm that always assigns the same score to all the atoms will achieve the best result, demonstrating weakness in ambiguity cover. The Average-distance evaluation is introduced and adopted as the main evaluation metric offering good balance between the proximity cover and ambiguity cover (**See Methods - Evaluation and SI Fig. SI-2**). Throughout the rest of this manuscript, “*Average-distance*” is referred to as “Evaluation Score”. Illustrative examples are provided in **SI Figure SI-5** to offer intuition and insight for different values of the evaluation score.

### ModiFinder Outperforms Baseline

We introduce three versions of the ModiFinder method. First, is the basic version of ModiFinder (MF-N). Second, is the refined version of ModiFinder (MF-R) that utilizes molecular formula filtration and substructure ambiguity refinement utilizing structurally related helper MS/MS spectra (**See Methods Substructure Refinement by Formula and Methods Refinement by Helpers**). Third, an Oracle Method (MF-O) is introduced that has knowledge of the true modification site to provide an approximate upper bound of ModiFinder performance. This is achieved by simulating the ability to reduce the ambiguity of substructure assignments to the MS/MS peaks **(See Methods Oracle)** by eliminating substructures that do not contain the modification site in shifted MS/MS peaks. Additionally, the site localization performance is evaluated on two random baselines: Random choice (RC) and Random Distribution (RD) (**See Methods Site Localization baselines and alternative approaches**). Finally, an alternative *in silico* prediction benchmarking approach is introduced which utilizes MS/MS fragmentation prediction of CFM-ID^20^ (**See Methods Site Localization baselines and alternative approaches**).

ModiFinder’s performance (MF-R in **Fig. 3-A** and **SI Fig. SI-4**) lies above the random baselines (RC and RD) and the CFM-ID approach and below the Oracle (MF-O). Specifically, MF-R when compared to RC and RD baselines shows an average Evaluation Score increase of 0.181 and 0.180, respectively, across all MS/MS pairs (**Fig. 3-A**). Moreover, MF-R outperforms these baselines in 81% and 80% percent of benchmark pairs (**SI Fig. SI-3 A**). In comparison, the MF-O version of ModiFinder outperforms the RC and RD in 85% of benchmark pairs (**SI Figure SI-3 B**) and shows an average increase of 0.208 and 0.207, respectively (**Fig. 3-A**). MF-R exhibits enhanced performance not only relative to random baseline comparisons but also over the CFM-ID based alternative approach evidenced by an increment of 0.266 in the average evaluation score (**Fig. 3-A**). Surprisingly, the CFM-ID based approach is found to be worse than RC and RD. This is because when simulating all regio-isomers, the resulting simulated MS/MS spectra were highly similar. This resulted in a nearly uniform likelihood distribution across all atoms, which was penalized in the “Ambiguity Cover” evaluation dimension. Because the “evaluation score” penalizes high levels of ambiguity, in other benchmark metrics that deemphasize “Ambiguity Cover”, e.g. “*Is-max*” (**See *SI Figure SI-2 and SI Note 1***).

**Figure 3.**
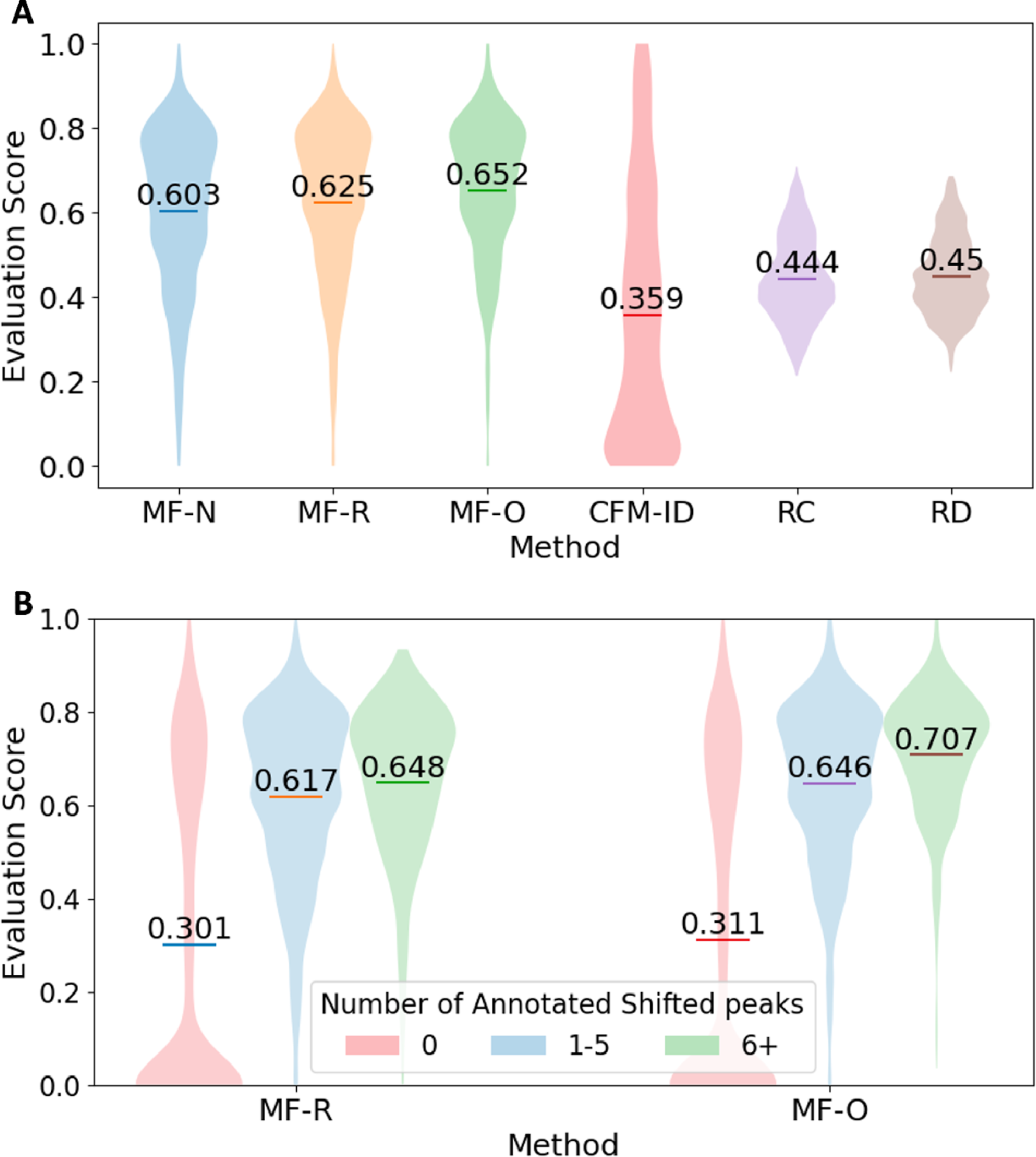
Performance Results. **A** Evaluation scores across pairs in all the libraries for different methods wher there is at least one shifted peak. ModiFinder outperforms the Random and CFM-ID baselines. MF-R, which utilizes helpers and formula constraints, improves upon MF-N, closing the gap to our upper bound performance of MF-O. **B** Evaluation score across pairs in all libraries for MF-O and MF-R based on the number of annotated shifted peaks. By increasing the number of shifted peaks, the performance of ModiFinder increases. This performance increase is consistent across different datasets and even for the MF-O demonstrating the utility of shifted peaks in finding modification sites.

We note that the gap in performance between MF-R and other baselines is greatest in lib4, which constitutes the majority of our database and might introduce a bias in the result. Nevertheless, MF-R maintains a clear advantage over the baseline across all the libraries (***SI Figure SI-4)***. Of the specific cases where MF-R does not outperform the RC and RD, manual analysis indicates in the majority of cases, this is because of a lack of shifted fragmentation peaks and or incomplete *in silico* substructure explanation of the MS/MS peaks.

With decreasing ambiguity of substructure assignments to MS/MS peaks, an increase in site localization performance is observed with MF-R improving upon MF-N, 0.625 vs 0.603 evaluation score respectively and MF-O, with the lowest ambiguity, further increases performance to 0.652 (**Fig. 3-A, Results Section Improving the Quality of Annotation and reducing the Ambiguity increases the Evaluation Score for more details)**.

### Importance of Peak Annotation

#### Shifted Peaks Matter in Site Localization

The results highlight the significance of annotated shifted peaks in the identification of the modification site where the MS/MS spectrum pairs featuring at least one shifted peak exhibit higher evaluation scores compared to those without any shifted peaks (**Fig. 3-B**). Furthermore, when there is an increase in the number of shifted peaks, the measured performance of MF-R and MF-O increases (**Fig. 3-B**). We hypothesize that the increase in shifted peaks potentially enriches the diversity of potential substructures, each MS/MS peak focusing on different segments of the compound. This diversity may reduce the overlap among substructures, thereby refining the precision in pinpointing the modification’s location. MF-O utilizes the shifted peaks more efficiently than the MF-R as it can remove the ambiguity introduced by more annotations on the extra shifted peaks.

#### Increasing the Fragmentation Depth Increases Ambiguity

One of the key steps of ModiFinder is substructure annotation of MS/MS fragment peaks. ModiFinder utilizes combinatorial fragmentation, i.e. MAGMa^16^ (one of the state of the art approaches^21,22^), to generate a set of potential fragmentation substructures for every MS/MS peak. Varying the fragmentation depth from two to four modulates the site localization performance. An increase in performance with MF-O is observed as the fragmentation depth increases (**Fig. 4-A**). This increase in performance can be attributed to the introduction of more substructures to the peaks, revealing the true substructure, especially for the peaks that were previously unannotated. Benchmark datasets show a 19% increase in the number of annotated shifted peaks when increasing fragmentation depth from two to four. While this increased explanation benefited MF-O, at higher fragmentation depths MF-R performance decreases. This is due to increased substructure ambiguity (average number of annotations assigned to each shifted peak). MF-O can counteract the increased ambiguity by utilizing the true fragmentation site to filter out substructures that do not benefit the localization. On the other hand, both the MF-R and MF-N are unable to filter incorrect substructures, leading to a loss of focus on the true modification site. Given the performance differences, fragmentation depth of 2 is chosen as the default value for MF-R. However, we anticipate that the optimal fragmentation depth might increase with enhancements in substructure assignments to MS/MS peaks.

**Figure 4.**
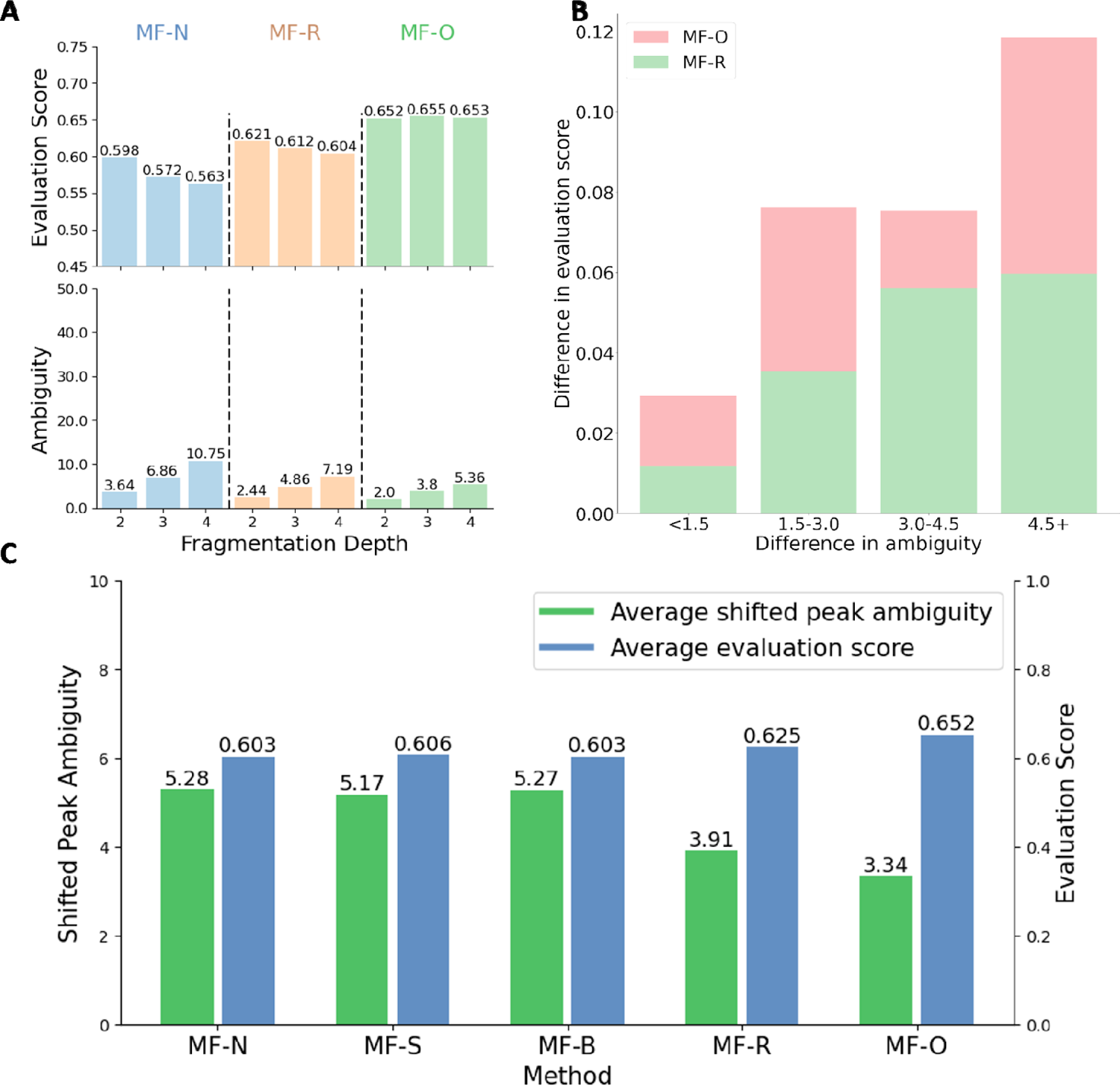
Correlation of ambiguity and performance. A) Impact of Fragmentation Depth on MF-N, MF-R, and MF-O Performance over pairs in all libraries. By increasing the fragmentation depth, the Oracle method (MF-O) attains higher scores, benefiting from more explanatory fragmentation. However, due to ambiguity introduced with more in-depth fragmentation, MF-N and MF-R do not benefit in site localization performance **B**) **Correlation of ambiguity and evaluation scores at the pairwise level over pairs in all libraries**. The evaluation score improvement from the unrefined ModiFinder (MF-N) for MF-O and MF-R based on the ambiguity reduction (difference in ambiguity). As the ambiguity difference increases, i.e., structural annotation becomes increasingly less ambiguous, the evaluation score difference increases. **C) Comparison of Average Ambiguity and Average Evaluation score across different settings of ModiFinder and Oracle for pairs in all datasets**. Methods that yield lower ambiguity correspondingly achieve higher evaluation scores.

#### Improving the Quality of Annotation and reducing the Ambiguity increases the Evaluation Score

MF-N calculates the likelihood scores using annotations derived from MAGMa^16^. Upon refining the molecular formula with SIRIUS^23^ (MF-S) or Buddy^24^ (MF-B) applied to MF-N, there was a decrease in ambiguity by 0.11 and 0.01, respectively (**Fig. 4-C**). This resulted in minimal improvements to the site localization evaluation score (**Fig. 4-C**). A larger magnitude reduction in MS/MS peak annotation ambiguity (reduction of 1.37) when helper compounds are utilized (MF-R) is observed. The most significant reduction in ambiguity (1.94) is observed, predictably, with the oracle (MF-O) (**Fig. 4-C**). There is a noticeable correlation between the decrease in annotation ambiguity and the improvement in evaluation scores; specifically, when the ambiguity of shifted peaks is reduced through refinements or the oracle’s knowledge, there is a corresponding increase in the evaluation score (**Fig. 4-C**). Additionally, the impact of ambiguity reduction was analyzed on a pair-by-pair basis. By categorizing pairs of MS/MS spectra based on the extent of ambiguity reduction, it was found that larger decreases in ambiguity corresponded to more significant improvements in site localization evaluation scores (**Fig. 4-B**). Figure **RSF-B** further indicates that for a comparable reduction in ambiguity, the oracle, on average, achieves greater improvements. This outcome is anticipated, as the oracle’s role extends beyond merely reducing ambiguity and skews the distribution of unambiguous peaks towards the actual modification site by eliminating substructures that do not contain the modification site.

#### Web User Interface for Domain-Based Improvements

An interactive analysis platform was developed to facilitate the utilization and refinement of site localization by chemists and mass spectrometrists using ModiFinder. Although the integration of helper compounds and formula refinement significantly enhances ModiFinder performance, these techniques do not fully eliminate substructure annotation ambiguity. Therefore, this interface enables expert users to apply their domain knowledge to eliminate incorrect substructures for each MS/MS peak. The web interface then synthesizes this user input (or multiple user refinements) with ModiFinder to produce a refined likelihood distribution.

As proof of principle, the web interface of ModiFinder and domain expertise were employed to solve the location of structural modifications of two natural products: *Kirromycin* and *Naphthomycin B*, two structurally complex natural product antibiotics^25,26^. These compounds were selected as they exhibited high structural complexity and there existed a large diversity of structural analogs that remain unidentified. First example demonstrates the ability to localize the N-methylation of *Kirromycin* that leads to its derivative *Goldinodox* (**Fig. 5**). In ModiFinder’s initial prediction, the true modification site was among the highest scoring. However, these existed two regions on the structure with non-zero likelihood scores (**Fig. 5-A**). MF-R computationally assigned seven possible substructures to each of the 112.04 *m/z* and 178.05 *m/z* peaks. By taking the polarization of the neighboring bonds to the carbonyl as well as the numbers of (single) bonds to be broken into account, domain experts were able to limit the substructures assigned to the peak 112.04 *m/z* to one (visualized by the green dotted rectangle in **Fig. 5**). Similarly, the set of substructures assigned to the 178.05 *m/z* peak was also manually reduced to one, i.e. the unlikely events of double bond breaking, forming of terminal amides through alkene loss, or alkene side chain methyl cleavage were eliminated. This reduction in ambiguity improved the site localization of the methylation and increased the evaluation score from 0.86 to 0.93.

**Figure 5.**
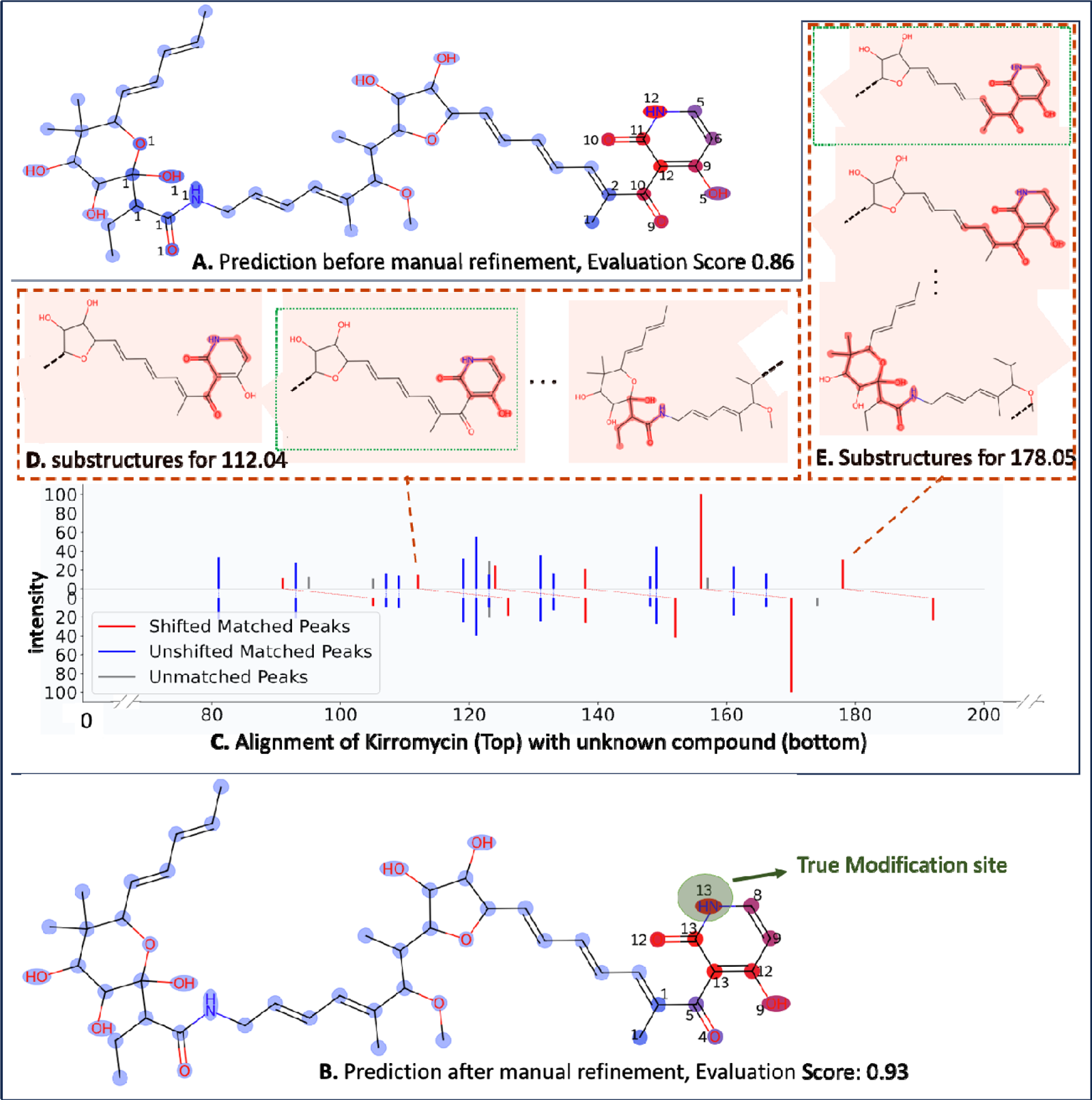
Combination of domain knowledge with ModiFinder’s user interface to improve the modification site prediction for Kirromycin. For the pair of *Kirromycin* and an unknown compound (*Goldinodox*), the initial prediction is improved by manually selecting the likely substructures for peak 112.04 m/z and 178.05 m/z improving the evaluation score from 0.86 to 0.93. **A** ModiFinder prediction before the refinement. **B** ModiFinder prediction after the refinement, the true modification site is highlighted by the green circle. **C** The alignment of *Kirromycin* and the unknown compound. The peaks of *Kirromycin* are shown at the top and the peaks of the unknown compound are shown at the bottom where shifted and unshifted peaks are highlighted with red and blue colors respectively. **D, E** shows the substructures assigned to peaks with 112.04 m/z and 178.05 m/z, due to the high number of substructures (ambiguity) only three substruc ures are shown. The green dotted box shows the substructure manually selected based on expert understanding of gas phase fragmentation.

Second example demonstrates the methylation of *Naphthomycin B* (**Fig. 6-A and B**). The specific challenge of *Naphthomycin B*, is due to the cyclic 2D structure, which causes MS/MS fragments (133.07 *m/z* and 147.08 *m/z*) to be ambiguous between multiple substructures around the 2D cycle (**Fig. 6-D**). This ambiguity leads to 24 high scoring sites (atoms) across the compound. In the case of *Naphthomycin B*, taking into account the likely gas phase fragmentation site at the amide bond, the substructure ambiguity for 133.07 *m/z* and 147.08 *m/z* decreased from 27 and 40 substructures to 2 substructures each. This resulted in a decrease of 24 high scoring modification sites to two high scoring modification sites above the true modification site. Further, the site localization was narrowed to a single likelihood region for the potential methyl-carrying site. This manual refinement improved the evaluation score from 0.56 to 0.74. However, ModiFinder, even given this ambiguity reduction, reported the highest likelihood two atoms away from the true site. The small number of shifted peaks likely limited the ability to localize to a specific site, but the ambiguity reduction provided by domain experts enabled ModiFinder to reach the limits of localization with the given MS/MS fragmentation.

**Figure 6.**
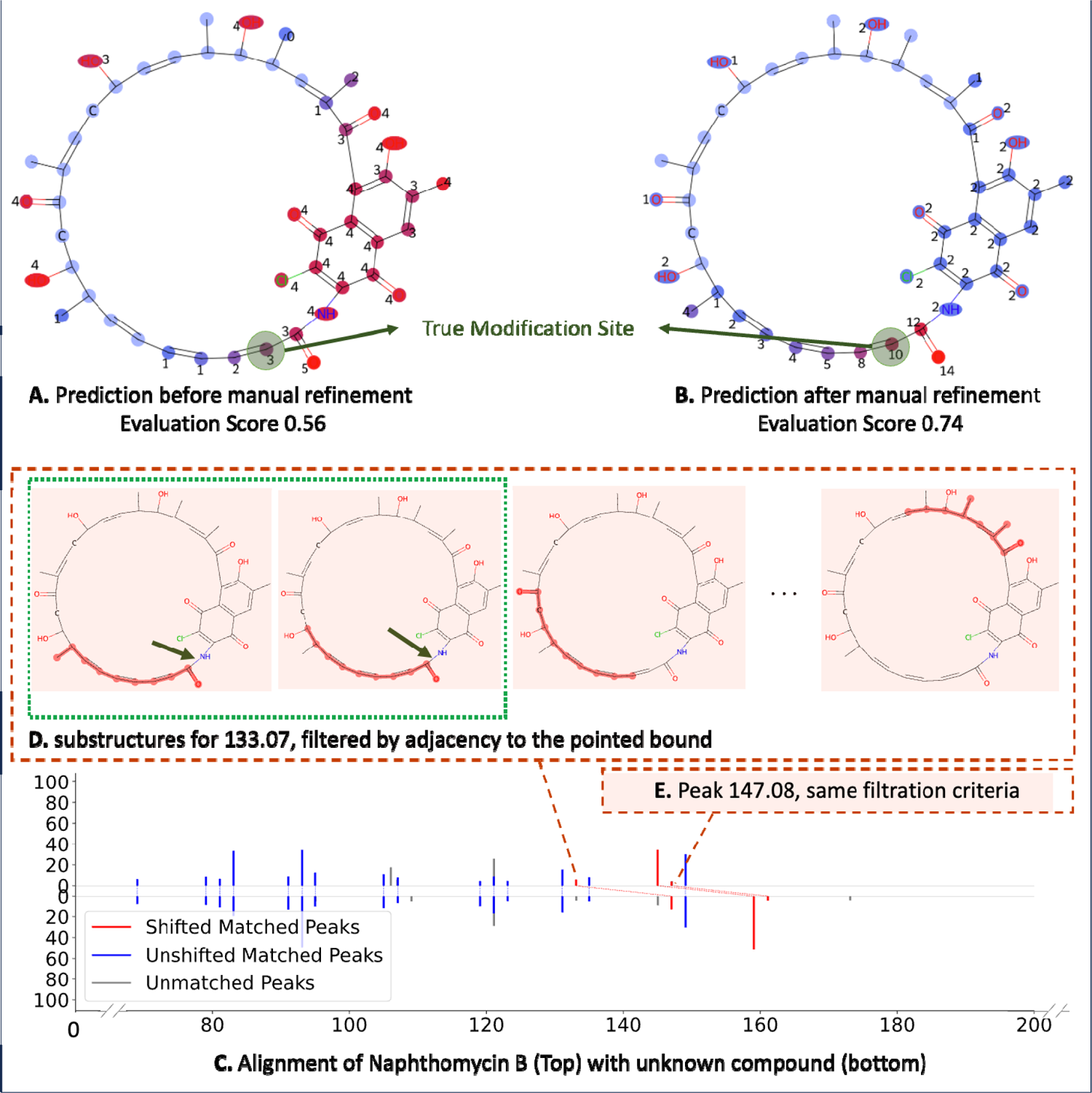
Combination of domain knowledge with ModiFinder’s user interface to improve the modification site prediction for Naphthomycin B. For the pair of *Naphthomycin B* and an unknown compound (*Naphthomycin A*), the initial prediction is improved by manually eliminating the unlikely substructures for peaks 133.07 m/z and 147.08 m/z, improving the evaluation score from 0.56 to 0.74. **A, B** ModiFinder prediction before and after the refinement, the modification site is highlighted by the green circle. **C** The alignment of *Naphthomycin B* and the unknown compound. The peaks of *Naphthomycin B* are shown at the top and the peaks of the unknown compound are shown at the bottom where shifted and unshifted peaks are highlighted with red and blue colors respectively. **D** shows the substructures as igned to peak with 133.07 m/z, due to the abundance of substructures (ambiguity) only four substructures are shown. The green dotted box shows the substructures manually selected based on expert understanding of gas phase fragmentation specific to amide bonds, which are indicated by green arrows. **E.** Same filtration applied to the peak with 147.08 m/z.

**Figure 7.**
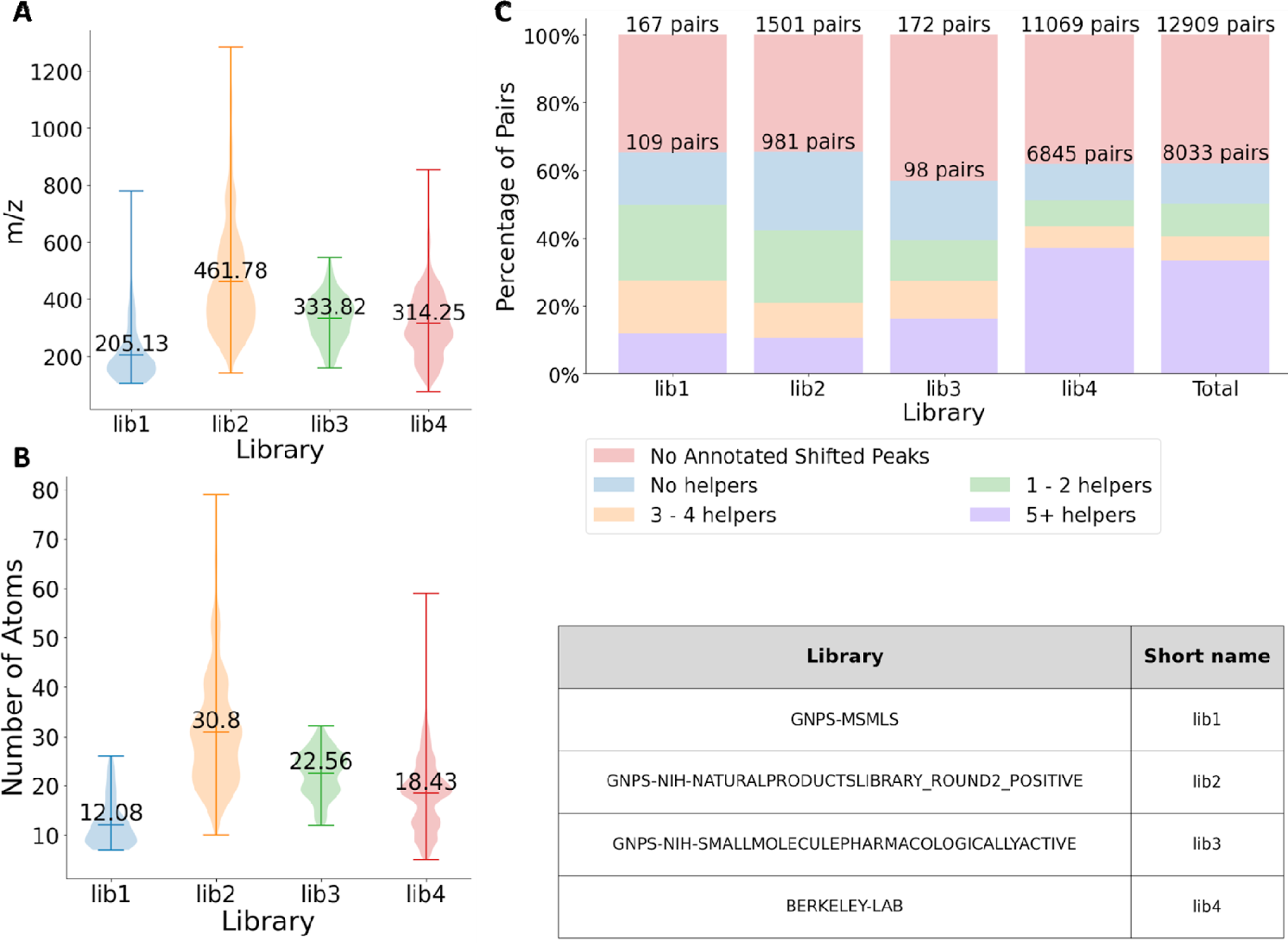
Distribution of Data in benchmarking libraries. **A** Shows the average and the distribution of m/z over the different libraries for pairs with at least one annotated shifted peak. **B** Shows the average and the distribution of the number of atoms in the compounds for each benchmark library for pairs with at least one annotated shifted peak. **C** For each library, the percentage of the pair of spectra with no annotated shifted peak is shown, the rest of the pairs are then categorized and shown based on their number of helpers. The majority of pairs have at least one shifted peak.

## Discussion

Here we introduced the challenge of site localization of chemical modifications in small molecules and presented our computational solution: ModiFinder. As demonstrated in our benchmarking results, ModiFinder and its refinements are able to outperform random baselines and *in silico* prediction alternative strategies. Promisingly, we also observe that due to refinements, ModiFinder makes significant progress to approach the performance of the oracle method, that is an estimate of the upper bound on performance. To bridge this gap for practical usage, our web interface enables an expert user to input their knowledge to bring the performance closer to the oracle performance and in some cases could even exceed the oracle.

Despite the promise of this presented work, we would reemphasize that the overall site localization problem as approached by ModiFinder is limited by two factors: first, the number of shifted peaks. Since the Oracle only simulates a reduction in substructure ambiguity, the Oracle’s performance is limited by the number of shifted peaks and the MS/MS fragmentation more generally. This is evidenced by the fact that our Oracle cannot reach perfect performance in our evaluation metric (**Fig. 3-A**) even with knowledge of the true modification site. Second, the substructure annotation ambiguity of the MS/MS peaks. Specifically, we note that ModiFinder and the Oracle method struggle in situations where the molecules exhibit high levels of 2D structure symmetry (**SI Note 2**). Given these limiting factors to ModiFinder performance, we anticipate (i) future instrumentation and method developments will produce richer and more complementary MS/MS fragmentation; (ii)computational and data acquisition advancements that aid in the reduction of substructure ambiguity of MS/MS fragmentation in small molecules, both of which will enhance ModiFinder’s performance going forward.

Although we have provided two example applications in the natural product field, we hope and anticipate that modification site localization will be broadly used in other communities that utilize small molecule untargeted mass spectrometry, e.g. toxicology, pharmacology, metabolism, exposomics, drug discovery, chemical biology, to name a few.

## Methods

### Definitions

MS2-known: The MS/MS spectra of known compound

S-known: The 2D chemical structure of the known compound

MS2-unknown : The MS/MS spectra of unknown compound

### ModiFinder Overview

Given the MS/MS of known (MS2-known) and unknown (MS2-unknown) compounds and the structure of the known compound (S-known) as input (**Fig. 2-A**), ModiFinder produces a likelihood distribution of the modification site location (**Fig. 2-D**). The process begins with ModiFinder assigning a set of potential substructures to the peaks in the MS2-known spectrum through *in-silico* fragmentation of S-known (**Fig. 2-B**). Subsequently, the MS2-known and MS2-unknown are aligned to find the matching peaks in each respective spectrum, producing peaks that have shifted in mass (shifted) and those that remain unchanged (unshifted). To predict the site localization, a likelihood score is assigned to each atom, where atoms in the substructures of the shifted peaks are rewarded while the atoms in the substructures in unshifted peaks are penalized (**Fig. 2-C**). Finally, The likelihood score of each atom is calculated. The following sections provide a detailed explanation of each step in this process.

### MS/MS Alignment

First, as a preprocessing step, all the peaks with intensities less than 1% of the base peak are removed, and the peaks are normalized to sum to a Euclidean norm of 1 to reduce noise^4^. Then, the GNPS^7^ alignment method is utilized to identify matched peaks between the known and unknown spectrum, by accounting for the mass delta of their respective precursors^7,27^. In the alignment process, the GNPS alignment method considers two types of matches: one where peaks have the same mass (non-shift), and another where peaks are offset by the difference in their precursor masses (shift). For each peak in the known compound’s spectrum, the availability of both non-shift and shift peaks are examined and all possible matched candidates of each peak are considered; Out of all these possibilities, the GNPS alignment method efficiently approximates the best-scoring match. Specifically, A bipartite graph is created where the nodes represent the peaks of MS2-known and MS2-unknown. An edge is drawn between an MS2-known peak and an MS2-unknown peak under two conditions: if their difference is less than a predefined threshold, indicating an unshifted match, or if it lies within the threshold range relative to the difference in precursor masses of the known and unknown compound, indicating an unshifted peak. The weight of each edge is the product of the intensities of the corresponding peaks. The goal is to find the maximum-scoring match. A greedy algorithm is used to approximate this matching. At each step, the maximum remaining edge is selected and added to the result. Then both ends of that edge along with all the edges connected to them are removed from the graph. In our experiments, a tolerance of 40 (ppm) is adopted as the threshold used to calculate the edges.

### Combinatorial Fragmentation and Refinement For Substructure Assignment

The peaks of MS2-known are annotated by assigning each peak a series of potential substructures using substructures generated from S-known following the MAGMa method^16^. In short, the fragmentation of S-known goes as follows: First, each of S-known’s heavy (non-hydrogen) atoms are removed once, each time yielding one or more substructures. Then the same process is repeated for each of the resulting substructures. The full fragmentation of the S-known is performed in a breadth-first search traversal. The generated substructures are stored as a bitstring where each bit represents one of the heavy atoms in S-known. ModiFinder begins with the initial structure (S-known) and continues the aforementioned breadth-first fragmentation approach up to a predetermined depth; a maximum depth of *2* is chosen here for the experimental setup.

Once the fragmentation step is done, for each substructure, the theoretical charged-m/z is calculated and compared to each peak’s m/z in the MS2-known. The maximum charge is assumed to be 1. If the theoretical m/z of a substructure falls within a specified m/z tolerance to the empirical m/z of a peak, then that substructure is assigned to that peak. Here, 40 ppm was chosen as the default m/z tolerance.

### Substructure Refinement by Formula

Predicted formulas provided by SIRIUS^23^ or BUDDY^24^ were used to filter the possible substructures for each MS/MS peak. SIRIUS, given the spectra of a compound, generates a pool of potential candidates using the information of the MS1. Next, it evaluates the interpretability of MS/MS spectra for each candidate by constructing a fragmentation tree^28^. The ModiFinder algorithm leverages this information by parsing the fragmentation tree and retrieving the formula assigned to each peak. This formula is then used to remove any substructure assigned to that peak with a different formula. Due to the performance complexity, SIRIUS is only computed for compounds with a precursor mass of 500 Da or less. For each compound, a mgf file is generated using the data retrieved from the MS/MS spectral library. This mgf file is then passed to v5.6.3 of the runnable script [https://github.com/boecker-lab/sirius/releases]. Then, the non-hydrogen part of the formula of each peak is compared with the formula of all the potential substructures assigned to that peak; filtering out all the substructures that have a different formula.

BUDDY’s ‘*assign_subformula*’ function is employed as an alternative to annotate formulas of fragmentation peaks^24^ with the same parameter set as ModiFinder (40 ppm is applied for the experiments), *-1.0* for the ‘*dbe_cutoff’* (the default value). The provided formulas are used to refine the substructures assigned to each peak by removing substructures that have different formulas.

### Refinement by Helpers

ModiFinder leverages additional compounds in MS/MS libraries that exhibit structural similarities to the known compound S-known, referring as helper compounds, to refine the substructure annotations. Specifically, the compounds within the same MS/MS library as S-known that share identical adducts and instruments but differ from S-known at precisely one modification site are identified as potential helper compounds. To ensure there is no information leakage and the unknown compound is not among the helpers, any compound that possesses a precursor mass within a 0.5 range of the precursor mass of the unknown compound is eliminated. Suppose H_S-known_ = {h_1_, …, h_n_} as the set of selected helper compounds for S-known. For each helper compound, denoted as h_i_, the same in-silico fragmentation process is performed on h_i_’s structure to annotate h_i_’s peaks. Next, h_i_’s spectra are aligned to the MS2-known to find matched peaks (shift and unshift). For every peak that has shifted, any substructure assigned to that peak in MS2-known that does not include the modification site between S-known and h_i_ is eliminated.

### Calculating the Site Localization

A score for each atom is computed, indicative of the likelihood of being the modification site. This score, termed as the “likelihood score” and shown by LJ (LJ_i_ for atom with index *i* in the graph), aims to serve as a score that measures the amount of evidence of an atom’s candidacy for being the site of modification. This scoring is performed under the assumption that there is only one modification site. Under this assumption, shifted peaks are probable hosts of the modification site. In Contrast, the atoms presenting in matched but unshifted peaks are penalized.

Each matched peak assigns a contribution score to each atom. i,j shows the contribution score assigned to atom j by peak i. For *i-*th matched peak, the scores for the atoms are calculated as follows: initially, the contribution scores for all atoms are assigned a value of 0. Then, assuming ***S*** _i_ *is the set of all the substructures assigned peak i:*

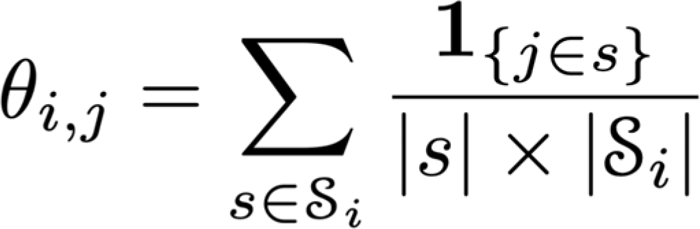

Where |s| is the number of atoms in substructure *s*, |***S*** _i_| is the number of structures assigned to peak i, and the is the indicator function that is *1* if j-th atom exists in substructure *s* and *0* otherwise.

Each matched peak itself receives a clarity score that represents how informative its substructures are. For example, if a peak has one substructure assigned to it but the substructure contains all the atoms in the compound or if the peak has multiple substructures assigned to it and overall each atom appears the same number of times, then the peak is not informative and must receive a low clarity score. Similarly, if the peak has few structures and they all focus on a specific and small part of the atom, then the peak is considered informative and will receive a high clarity score. To compute this clarity score, the Shannon entropy^29^ is calculated. The clarity score of the peak *i, C_i_,* is proportional to this entropy score:

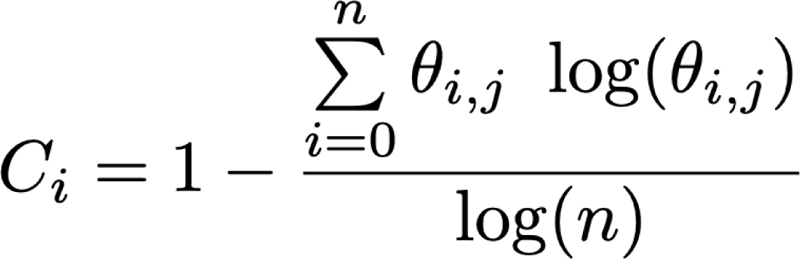

Where n is the total number of atoms. Finally, □_j_ is updated. If the peak is shifted, it is increased by c_i_ x □_i,j_, and if it is unshifted, □_j_ is decreased by c_i_ x □_i,j_.

For the final step, after normalizing □ so that the maximum value is 1, Then any value below 0.5 is set to 0 and to further highlight the differences, especially in the high-scoring atoms, all the values are raised to the power of 4 as a dynamic range adjustment. Finally, the values are normalized again to have a sum of 1.

### Evaluation Score

To measure the performance of ModiFinder and compare it to alternative approaches and baselines, an evaluation function is needed. This evaluation function takes in the predicted likelihood array together along with the true modification site, i.e. the 2D graph structure of the known compound and the actual modification site location. The evaluation produces a score between zero and one. Scores approaching one signify more accurate predictions, while those closer to zero indicate less accurate predictions.

The Average-distance evaluation method is proposed for the evaluation.

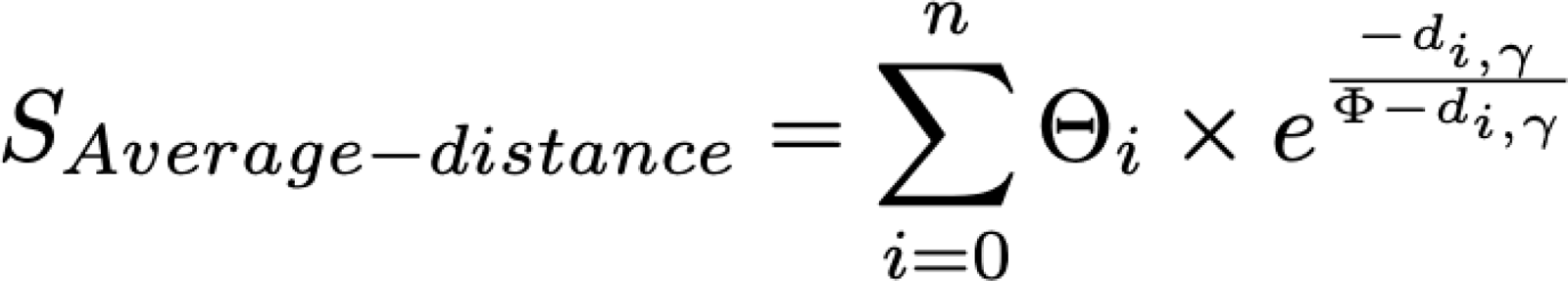

Where d_i,__ denotes the distance between the atom with index ‘i’ and true modification site □ on the 2D graph structure and denotes the diameter (greatest shortest distance between any two nodes in the grap) of the 2D graph structure. Using the diameter helps normalize the distances based on the size and structure of the compound. Normalization ensures uniformity in the evaluation metric across molecules of varying sizes. In Average-distance, the impact of each atom on the total score decreases exponentially with its distance from the actual modification site. Atoms with high predicted likelihood situated far away from the true site contribute less to the evaluation score, whereas those in closer proximity contribute more. This aspect addresses the Average-distance’s capacity to account for the proximity cover. In addition, since the scores are norm lized to have a sum of 1, the likelihood scores are directly proportional to each atom’s relative influence. Consequently, in ambiguous scenarios where many atoms have high predicted likelihood, the relative likelihood of a single atom diminished. This reduction, in turn, lessens their overall effect on the evaluation score, encapsulates the method’s ambiguity cover (**SI Figure SI-5**). Beyond this evaluation score, the performance of ModiFinder is also examined over *Is-max*, *Proximity*, and *Sorted-rank* evaluation methods (See **SI Note 1**).

### Site Localization baselines and alternative approaches

Random Choice (RC) adopts a random selection approach for designating one of the atoms as the modification site. In contrast, our second baseline, termed “Random Distribution” assigns a likelihood score to atom *i*, where The Oracle approach is built on top of ModiFinder and uses the extra information of the true modification site. After ModiFinder has annotated the peaks with putative substructures using the combinatorial fragmentation and formula and helper refinements, Oracle applies an extra elimination step. Specifically, for every shifted peak of MS2-known, Oracle filters out any substructure assigned to that peak that does not contain the true modification site. Once this step is completed, all the substructures assigned to the shifted peaks are guaranteed to contain the modification site.

In addition, utlizing the MS/MS fragmentation prediction property of CFM-ID^20^ a different modification site method is developed as an alternative approach and a baseline to compare against. CFM-ID is a tool to predict spectra based on a given molecular structure. This alternative approach is only developed for evaluation as it uses the structure of the “unknown compound” which is paradoxical and impractical in real-world scenarios. It serves merely as a reference for comparison, and the ability of ModiFinder to surpass its performance, despite the latter’s theoretical omniscience, further emphasizes the effectiveness of ModiFinder. This alternative method is designed to use CFM-ID as a black box to find the modification site (Refer to SI **Fig. SI-6** for a visual illustration of this method). First, using the extra information provided by the structure of the “unknown” compound, the modification substructure is calculated. Then, this modification substructure is permuted across the known structure (S-known). With each permutation, the modification is attached to an atom in S-known, creating an analog to S-known and a possible candidate for the unknown compound. To attach the modified part to an atom, first the same original bond type is tried, if that does not produce a valid structure, other bond types are tried. After this step, CFM-ID tool is used to predict spectra for each structure (SI **Figure SI-6. C.**). To run CFM-ID, the docker container provided [https://hub.docker.com/r/wishartlab/cfmid is used with 0.001 for ‘prob_thresh’ (the default value), ‘trained_models_cfmid4.0/[M+H]+/param_output.log’ for param_file, ‘/trained_models_cfmid4.0/[M+H]+/param_config.txt’ for config file.

The similarity of the predicted spectra and MS2-unknown is measured using the cosine similarity score (**SI Figure SI-6. D)**. This similarity is reported as the likelihood score of the atom corresponding to the permutation SI (**Figure SI-6. E.)**. In addition to the visualization of the algorithm, two examples for Deoxyadenosine and Deoxyadenosine Monophosphate (**Fig. SI-7**), and Tyramine and 3-MethoxyTyramine (**Fig. SI-8**) are also provided.

### MS/MS Spectral Library Data Preparation

The creation of the database used for ModiFinder evaluation involved a multi-step process. Initially, compounds with known 2D structures were selected from MS/MS libraries containing compounds with known structure^30–35^. the following public MS/MS spectral libraries were used to retrieve the MS/MS and structure pairs: [1-GNPS-MSMLS, 2-GNPS-NIH-NATURALPRODUCTSLIBRARY_ROUND2_POSITIVE, 3-GNPS-NIH-SMALLMOLECULEPHARMACOLOGICALLYACTIVE^34^, and 4-BERKELEY-LAB]. In addition, data from the TUEBINGEN-NATURAL-PRODUCT-COLLECTION was used for the web tool performance demonstration.

For each library, every possible pair in that library are analyzed to verify their eligibility. For each pair with known structure, (i) their precursor mass is checked to be less than 2000 Da, (ii) the difference in precursor masses is less than 50% of the precursor mass of the smaller compound, (iii) they share the same *M+H* adduct, and finally, (iv) the structures are examined to differ in exactly once modification site. In the final verification step, both SMILES structures are converted to an RDKit^36^ molecule object, then the “*GetSubstructMatch”* function is called on the heavier compound’s object with the smaller compound’s object as input. If the smaller compound is a substructure of the larger compound, the number of edges between the atoms in the substructure set and the atoms not in the substructure set is calculated as the number of modification sites. Any pair with more than one edge is discarded.

### Natural Product Data Acquisition Methods

*Kirromycin*, *Goldinodox*, *Naphthomycin B*, and *Naphthomycin A* samples were dissolved in DMSO. Subsequently, the samples were pooled and diluted with ACN/water (80/20) to a final concentration of 8 µg/mL (Kirromycin, Goldinodox) resp. 7 µg/mL (Naphthomycin A and B). LC-MS/MS Data was acquired on a Vanquish ultrahigh-performance liquid chromatography (UHPLC) coupled to a Q executive HF apparatus (Thermo Fisher Scientific) equipped with an heated electrospray ionization (HESI) source. The chromatographic separation was performed with a constant flow rate of 0.5 mL/min with mobile phase A (H_2_O +0.1% formic acid) and mobile phase B (ACN +0.1% formic acid). The separation gradient started with 5% B as initial conditions, which was linearly increased to 50% B at 8 minutes, and then to 99% B at 10 minutes, followed by a washing phase with 99% B, and a re-equilibration phase at the initial conditions. As reversed phase HPLC column, a Phenomenex Kinetex 1.7 µm EVO C18 RP, 100 Å pore size, dimension: 50 x 2.0 (mm, length x inner diameter) was employed. The MS/MS method was previously optimized^37^, the HESI source parameters were set as follows: auxiliary gas flow and temperature were respectively 12 AU and 400 °C, sweep gas flow was 1 L/min, sheath gas flow rate was set to 50 AU. The MS was operated in positive mode, The spray voltage was set to 3.5 kV while an inlet capillary temperature of 250 °C was adopted. The scan range was set to 150-1500 m/z and the resolution to 30,000. The fragmentation was performed in Data Dependent Analysis (DDA) mode, the 5 most abundant ions were fragmented per MS survey scan with a resolution of 15,000, an isolation window of 1 m/z, and with the following stepped normalized collision energy: 25, 35, 45 eV. Finally, data were deposited as a spectral library in GNPS (https://external.gnps2.org/gnpslibrary/TUEBINGEN-NATURAL-PRODUCT-COLLECTION.mgf).

## Code Availability

The standalone source code of the ModiFinder tool (written in python) can be accessed via https://github.com/mshahneh/SmallMol_Mod_Site_Localization

The web-interface software for ModiFinder can be found here: https://github.com/mshahneh/ModiFinder-dash

Additionally, all the necessary code to replicate the experiments, benchmarking, and analysis conducted with ModiFinder is provided at: https://github.com/mshahneh/ModiFinder_Analysis

A public and interactive version of ModiFinder can be accessed here: https://modifinder.gnps2.org/

Please note that this website is continually improved and results may not match exactly as presented in this manuscript. To reproduce the results, please find the software deposited in Zenodo.

The specific version of the repositories utilized in this study has been archived on Zenodo here:

XXX

XXX

XXX

## Data Availability

The benchmarking libraries for ModiFinder are available for download at https://external.gnps2.org/gnpslibrary.

The set of pairs with single modifications for benchmarking in this manuscript are available here:

https://www.dropbox.com/scl/fi/t07mpuzdde2k3oe0hh15o/GNPS-MSMLS.csv?rlkey=xh98imxmri5b82y3lipwsfd39&dl=0 https://www.dropbox.com/scl/fi/8nw80ayxshk2lpti6mr4r/BERKELEY-LAB.csv?rlkey=udv2rjxbk9ol02y6gx5kc4lwg&dl=0 https://www.dropbox.com/scl/fi/ni88vniggcy21eif0v3c8/GNPS-NIH-NATURALPRODUCTSLIBRARY_ROUND2_POSITIVE.csv?rlkey=y18q7bea8i64rgz96cb174ovc&dl=0 https://www.dropbox.com/scl/fi/8nw80ayxshk2lpti6mr4r/BERKELEY-LAB.csv?rlkey=udv2rjxbk9ol02y6gx5kc4lwg&dl=0

The CFM-ID Results are available here:

https://www.dropbox.com/scl/fi/hcg76wkqa0r8lj9p0y3dp/cfmid_exp.zip?rlkey=371yuhokh20l5btpeklq7vqh7&dl=0

The pairs of helpers we used for refinement of sub-structure annotation can be found here: https://www.dropbox.com/scl/fi/55r9cyjanw9cft95v65x5/helpers.zip?rlkey=vm3jltxbnq2f4c998chgs7m6h&dl=0

The Sirius computed results can be found here: https://www.dropbox.com/scl/fi/ejnfd0j04qrjzryj5i7cl/SIRIUS.zip?rlkey=bxnyxefol5tfojp4cdmsyl8d2&dl=0

## Supplementary information

https://docs.google.com/document/d/1zjkrOw1CWtgu0b-tQYr4HDKcyq3ukH_82OJdDp2CBlQ/edit?usp=sharing

## Acknowledgments

MW and MRZS are supported by NIH 5U24DK133658-02 and an Agilent Inc. Gift. DP was supported through the German Research Foundation, through the CMFI Cluster of Excellence (EXC 2124) and the Collaborative Research Center CellMap (TRR 261). We thank Prof. YE was supported by NIH 1R03OD034493-01. We thank Heike Broetz-Oesterhelt for providing natural product standards. NG was supported by the National Science Foundation (NSF) CAREER award (Award No. 2047235). ATA was supported by University of Denver start-up funds.VVP was supported by NIH R35GM128690 and the ALSAM Foundation.

## Bibliography

(1) Schrimpe-Rutledge, A. C.; Codreanu, S. G.; Sherrod, S. D.; McLean, J. A. Untargeted Metabolomics Strategies-Challenges and Emerging Directions. J. Am. Soc. Mass Spectrom. 2016, 27 (12), 1897–1905. 10.1007/s13361-016-1469-y.

(2) McLuckey, S. A. Principles of Collisional Activation in Analytical Mass Spectrometry. J. Am. Soc. Mass Spectrom. 1992, 3 (6), 599–614. 10.1016/1044-0305(92)85001-Z.

(3) Stein, S. E.; Scott, D. R. Optimization and Testing of Mass Spectral Library Search Algorithms for Compound Identification. J. Am. Soc. Mass Spectrom. 1994, 5 (9), 859–866. 10.1016/1044-0305(94)87009-8.

(4) Bittremieux, W.; Wang, M.; Dorrestein, P. C. The Critical Role That Spectral Libraries Play in Capturing the Metabolomics Community Knowledge. Metabolomics Off. J. Metabolomic Soc. 2022, 18 (12), 94. 10.1007/s11306-022-01947-y.

(5) Cooper, B. T.; Yan, X.; Simón-Manso, Y.; Tchekhovskoi, D. V.; Mirokhin, Y. A.; Stein, S. E. Hybrid Search: A Method for Identifying Metabolites Absent from Tandem Mass Spectrometry Libraries. Anal. Chem. 2019, 91 (21), 13924–13932. 10.1021/acs.analchem.9b03415.

(6) Yang, J. Y.; Sanchez, L. M.; Rath, C. M.; Liu, X.; Boudreau, P. D.; Bruns, N.; Glukhov, E.; Wodtke, A.; de Felicio, R.; Fenner, A.; Wong, W. R.; Linington, R. G.; Zhang, L.; Debonsi, H. M.; Gerwick, W. H.; Dorrestein, P. C. Molecular Networking as a Dereplication Strategy. J. Nat. Prod. 2013, 76 (9), 1686– 1699. 10.1021/np400413s.

(7) Wang, M.; Carver, J. J.; Phelan, V. V.; Sanchez, L. M.; Garg, N.; Peng, Y.; Nguyen, D. D.; Watrous, J.; Kapono, C. A.; Luzzatto-Knaan, T.; Porto, C.; Bouslimani, A.; Melnik, A. V.; Meehan, M. J.; Liu, W.-T.; Crüsemann, M.; Boudreau, P. D.; Esquenazi, E.; Sandoval-Calderón, M.; Kersten, R. D.; Pace, L. A.; Quinn, R. A.; Duncan, K. R.; Hsu, C.-C.; Floros, D. J.; Gavilan, R. G.; Kleigrewe, K.; Northen, T.; Dutton, R. J.; Parrot, D.; Carlson, E. E.; Aigle, B.; Michelsen, C. F.; Jelsbak, L.; Sohlenkamp, C.; Pevzner, P.; Edlund, A.; McLean, J.; Piel, J.; Murphy, B. T.; Gerwick, L.; Liaw, C.-C.; Yang, Y.-L.; Humpf, H.-U.; Maansson, M.; Keyzers, R. A.; Sims, A. C.; Johnson, A. R.; Sidebottom, A. M.; Sedio, B. E.; Klitgaard, A.; Larson, C. B.; P, C. A. B.; Torres-Mendoza, D.; Gonzalez, D. J.; Silva, D. B.; Marques, L. M.; Demarque, D. P.; Pociute, E.; O’Neill, E. C.; Briand, E.; Helfrich, E. J. N.; Granatosky, E. A.; Glukhov, E.; Ryffel, F.; Houson, H.; Mohimani, H.; Kharbush, J. J.; Zeng, Y.; Vorholt, J. A.; Kurita, K. L.; Charusanti, P.; McPhail, K. L.; Nielsen, K. F.; Vuong, L.; Elfeki, M.; Traxler, M. F.; Engene, N.; Koyama, N.; Vining, O. B.; Baric, R.; Silva, R. R.; Mascuch, S. J.; Tomasi, S.; Jenkins, S.; Macherla, V.; Hoffman, T.; Agarwal, V.; Williams, P. G.; Dai, J.; Neupane, R.; Gurr, J.; Rodríguez, A. M. C.; Lamsa, A.; Zhang, C.; Dorrestein, K.; Duggan, B. M.; Almaliti, J.; Allard, P.-M.; Phapale, P.; Nothias, L.-F.; Alexandrov, T.; Litaudon, M.; Wolfender, J.-L.; Kyle, J. E.; Metz, T. O.; Peryea, T.; Nguyen, D.-T.; VanLeer, D.; Shinn, P.; Jadhav, A.; Müller, R.; Waters, K. M.; Shi, W.; Liu, X.; Zhang, L.; Knight, R.; Jensen, P. R.; Palsson, B. O.; Pogliano, K.; Linington, R. G.; Gutiérrez, M.; Lopes, N. P.; Gerwick, W. H.; Moore, B. S.; Dorrestein, P. C.; Bandeira, N. Sharing and Community Curation of Mass Spectrometry Data with Global Natural Products Social Molecular Networking. Nat. Biotechnol. 2016, 34 (8), 828–837. 10.1038/nbt.3597.

(8) Xing, S.; Hu, Y.; Yin, Z.; Liu, M.; Tang, X.; Fang, M.; Huan, T. Retrieving and Utilizing Hypothetical Neutral Losses from Tandem Mass Spectra for Spectral Similarity Analysis and Unknown Metabolite Annotation. Anal. Chem. 2020, 92 (21), 14476–14483. 10.1021/acs.analchem.0c02521.

(9) Watrous, J.; Roach, P.; Alexandrov, T.; Heath, B. S.; Yang, J. Y.; Kersten, R. D.; van der Voort, M.; Pogliano, K.; Gross, H.; Raaijmakers, J. M.; Moore, B. S.; Laskin, J.; Bandeira, N.; Dorrestein, P. C. Mass Spectral Molecular Networking of Living Microbial Colonies. Proc. Natl. Acad. Sci. U. S. A. 2012, 109 (26), E1743–1752. 10.1073/pnas.1203689109.

(10) da Silva, R. R.; Wang, M.; Nothias, L.-F.; van der Hooft, J. J. J.; Caraballo-Rodríguez, A. M.; Fox, E.; Balunas, M. J.; Klassen, J. L.; Lopes, N. P.; Dorrestein, P. C. Propagating Annotations of Molecular Networks Using in Silico Fragmentation. PLoS Comput. Biol. 2018, 14 (4), e1006089. 10.1371/journal.pcbi.1006089.

(11) Doll, S.; Burlingame, A. L. Mass Spectrometry-Based Detection and Assignment of Protein Posttranslational Modifications. ACS Chem. Biol. 2015, 10 (1), 63–71. 10.1021/cb500904b.

(12) Yu, F.; Teo, G. C.; Kong, A. T.; Haynes, S. E.; Avtonomov, D. M.; Geiszler, D. J.; Nesvizhskii, A. I. Identification of Modified Peptides Using Localization-Aware Open Search. Nat. Commun. 2020, 11 (1), 4065. 10.1038/s41467-020-17921-y.

(13) Chalkley, R. J.; Clauser, K. R. Modification Site Localization Scoring: Strategies and Performance. Mol. Cell. Proteomics MCP 2012, 11 (5), 3–14. 10.1074/mcp.R111.015305.

(14) Ruttkies, C.; Neumann, S.; Posch, S. Improving MetFrag with Statistical Learning of Fragment Annotations. BMC Bioinformatics 2019, 20 (1), 376. 10.1186/s12859-019-2954-7.

(15) Wolf, S.; Schmidt, S.; Müller-Hannemann, M.; Neumann, S. In Silico Fragmentation for Computer Assisted Identification of Metabolite Mass Spectra. BMC Bioinformatics 2010, 11, 148. 10.1186/1471-2105-11-148.

(16) Ridder, L.; Van Der Hooft, J. J. J.; Verhoeven, S.; De Vos, R. C. H.; Van Schaik, R.; Vervoort, J. Substructure-based Annotation of High resolution Multistage MS *^n^* Spectral Trees. Rapid Commun. Mass Spectrom. 2012, 26 (20), 2461–2471. 10.1002/rcm.6364.

(17) Wang, Y.; Kora, G.; Bowen, B. P.; Pan, C. MIDAS: A Database-Searching Algorithm for Metabolite Identification in Metabolomics. Anal. Chem. 2014, 86 (19), 9496–9503. 10.1021/ac5014783.

(18) Tsugawa, H.; Kind, T.; Nakabayashi, R.; Yukihira, D.; Tanaka, W.; Cajka, T.; Saito, K.; Fiehn, O.; Arita, M. Hydrogen Rearrangement Rules: Computational MS/MS Fragmentation and Structure Elucidation Using MS-FINDER Software. Anal. Chem. 2016, 88 (16), 7946–7958. 10.1021/acs.analchem.6b00770.

(19) Van Tetering, L.; Spies, S.; Wildeman, Q. D. K.; Houthuijs, K. J.; Van Outersterp, R. E.; Martens, J.; Wevers, R. A.; Wishart, D. S.; Berden, G.; Oomens, J. A Spectroscopic Test Suggests That Fragment Ion Structure Annotations in MS/MS Libraries Are Frequently Incorrect. Commun. Chem. 2024, 7 (1), 30. 10.1038/s42004-024-01112-7.

(20) Wang, F.; Liigand, J.; Tian, S.; Arndt, D.; Greiner, R.; Wishart, D. S. CFM-ID 4.0: More Accurate ESI-MS/MS Spectral Prediction and Compound Identification. Anal. Chem. 2021, 93 (34), 11692–11700. 10.1021/acs.analchem.1c01465.

(21) Blaženović, I.; Kind, T.; Torbašinović, H.; Obrenović, S.; Mehta, S. S.; Tsugawa, H.; Wermuth, T.; Schauer, N.; Jahn, M.; Biedendieck, R.; Jahn, D.; Fiehn, O. Comprehensive Comparison of in Silico MS/MS Fragmentation Tools of the CASMI Contest: Database Boosting Is Needed to Achieve 93% Accuracy. J. Cheminformatics 2017, 9 (1), 32. 10.1186/s13321-017-0219-x.

(22) Verdegem, D.; Lambrechts, D.; Carmeliet, P.; Ghesquière, B. Improved Metabolite Identification with MIDAS and MAGMa through MS/MS Spectral Dataset-Driven Parameter Optimization. Metabolomics 2016, 12 (6), 98. 10.1007/s11306-016-1036-3.

(23) Dührkop, K.; Fleischauer, M.; Ludwig, M.; Aksenov, A. A.; Melnik, A. V.; Meusel, M.; Dorrestein, P. C.; Rousu, J.; Böcker, S. SIRIUS 4: A Rapid Tool for Turning Tandem Mass Spectra into Metabolite Structure Information. Nat. Methods 2019, 16 (4), 299–302. 10.1038/s41592-019-0344-8.

(24) Xing, S.; Shen, S.; Xu, B.; Li, X.; Huan, T. BUDDY: Molecular Formula Discovery via Bottom-up MS/MS Interrogation. Nat. Methods 2023, 20 (6), 881–890. 10.1038/s41592-023-01850-x.

(25) Keller-Schierlein, W.; Meyer, M.; Zeeck, A.; Damberg, M.; Machinek, R.; Zähner, H.; Lazar, G. Isolation and Structural Elucidation of Naphthomycins B and C. J. Antibiot. (Tokyo) 1983, 36 (5), 484–492. 10.7164/antibiotics.36.484.

(26) Wolf, H.; Chinali, G.; Parmeggiani, A. Kirromycin, an Inhibitor of Protein Biosynthesis That Acts on Elongation Factor Tu. Proc. Natl. Acad. Sci. 1974, 71 (12), 4910–4914. 10.1073/pnas.71.12.4910.

(27) Bittremieux, W.; Schmid, R.; Huber, F.; van der Hooft, J. J. J.; Wang, M.; Dorrestein, P. C. Comparison of Cosine, Modified Cosine, and Neutral Loss Based Spectrum Alignment For Discovery of Structurally Related Molecules. J. Am. Soc. Mass Spectrom. 2022, 33 (9), 1733–1744. 10.1021/jasms.2c00153.

(28) Rasche, F.; Svatos, A.; Maddula, R. K.; Böttcher, C.; Böcker, S. Computing Fragmentation Trees from Tandem Mass Spectrometry Data. Anal. Chem. 2011, 83 (4), 1243–1251. 10.1021/ac101825k.

(29) Shannon, C. E. A Mathematical Theory of Communication. Bell Syst. Tech. J. 1948, 27 (3), 379–423. 10.1002/j.1538-7305.1948.tb01338.x.

(30) Petras, D.; Phelan, V. V.; Acharya, D.; Allen, A. E.; Aron, A. T.; Bandeira, N.; Bowen, B. P.; Belle-Oudry, D.; Boecker, S.; Cummings, D. A.; Deutsch, J. M.; Fahy, E.; Garg, N.; Gregor, R.; Handelsman, J.; Navarro-Hoyos, M.; Jarmusch, A. K.; Jarmusch, S. A.; Louie, K.; Maloney, K. N.; Marty, M. T.; Meijler, M. M.; Mizrahi, I.; Neve, R. L.; Northen, T. R.; Molina-Santiago, C.; Panitchpakdi, M.; Pullman, B.; Puri, A. W.; Schmid, R.; Subramaniam, S.; Thukral, M.; Vasquez-Castro, F.; Dorrestein, P. C.; Wang, M. GNPS Dashboard: Collaborative Exploration of Mass Spectrometry Data in the Web Browser. Nat. Methods 2022, 19 (2), 134–136. 10.1038/s41592-021-01339-5.

(31) Tobias, N. J.; Wolff, H.; Djahanschiri, B.; Grundmann, F.; Kronenwerth, M.; Shi, Y.-M.; Simonyi, S.; Grün, P.; Shapiro-Ilan, D.; Pidot, S. J.; Stinear, T. P.; Ebersberger, I.; Bode, H. B. Natural Product Diversity Associated with the Nematode Symbionts Photorhabdus and Xenorhabdus. Nat. Microbiol. 2017, 2 (12), 1676–1685. 10.1038/s41564-017-0039-9.

(32) Nothias, L.-F.; Petras, D.; Schmid, R.; Dührkop, K.; Rainer, J.; Sarvepalli, A.; Protsyuk, I.; Ernst, M.; Tsugawa, H.; Fleischauer, M.; Aicheler, F.; Aksenov, A. A.; Alka, O.; Allard, P.-M.; Barsch, A.; Cachet, X.; Caraballo-Rodriguez, A. M.; Da Silva, R. R.; Dang, T.; Garg, N.; Gauglitz, J. M.; Gurevich, A.; Isaac, G.; Jarmusch, A. K.; Kameník, Z.; Kang, K. B.; Kessler, N.; Koester, I.; Korf, A.; Le Gouellec, A.; Ludwig, M.; Martin H., C.; McCall, L.-I.; McSayles, J.; Meyer, S. W.; Mohimani, H.; Morsy, M.; Moyne, O.; Neumann, S.; Neuweger, H.; Nguyen, N. H.; Nothias-Esposito, M.; Paolini, J.; Phelan, V. V.; Pluskal, T.; Quinn, R. A.; Rogers, S.; Shrestha, B.; Tripathi, A.; Van Der Hooft, J. J. J.; Vargas, F.; Weldon, K. C.; Witting, M.; Yang, H.; Zhang, Z.; Zubeil, F.; Kohlbacher, O.; Böcker, S.; Alexandrov, T.; Bandeira, N.; Wang, M.; Dorrestein, P. C. Feature-Based Molecular Networking in the GNPS Analysis Environment. Nat. Methods 2020, 17 (9), 905–908. 10.1038/s41592-020-0933-6.

(33) Fox Ramos, A. E.; Le Pogam, P.; Fox Alcover, C.; Otogo N’Nang, E.; Cauchie, G.; Hazni, H.; Awang, K.; Bréard, D.; Echavarren, A. M.; Frédérich, M.; Gaslonde, T.; Girardot, M.; Grougnet, R.; Kirillova, M. S.; Kritsanida, M.; Lémus, C.; Le Ray, A.-M.; Lewin, G.; Litaudon, M.; Mambu, L.; Michel, S.; Miloserdov, F. M.; Muratore, M. E.; Richomme-Peniguel, P.; Roussi, F.; Evanno, L.; Poupon, E.; Champy, P.; Beniddir, M.A. Collected Mass Spectrometry Data on Monoterpene Indole Alkaloids from Natural Product Chemistry Research. Sci. Data 2019, 6 (1), 15. 10.1038/s41597-019-0028-3.

(34) Gauglitz, J. M.; Aceves, C. M.; Aksenov, A. A.; Aleti, G.; Almaliti, J.; Bouslimani, A.; Brown, E. A.; Campeau, A.; Caraballo-Rodríguez, A. M.; Chaar, R.; Da Silva, R. R.; Demko, A. M.; Di Ottavio, F.; Elijah, E.; Ernst, M.; Ferguson, L. P.; Holmes, X.; Jarmusch, A. K.; Jiang, L.; Kang, K. B.; Koester, I.; Kwan, B.; Li, J.; Li, Y.; Melnik, A. V.; Molina-Santiago, C.; Ni, B.; Oom, A. L.; Panitchpakdi, M. W.; Petras, D.; Quinn, R.; Sikora, N.; Spengler, K.; Teke, B.; Tripathi, A.; Ul-Hasan, S.; Van Der Hooft, J. J. J.; Vargas, F.; Vrbanac, A.; Vu, A. Q.; Wang, S. C.; Weldon, K.; Wilson, K.; Wozniak, J. M.; Yoon, M.; Bandeira, N.; Dorrestein, P. C. Untargeted Mass Spectrometry-Based Metabolomics Approach Unveils Molecular Changes in Raw and Processed Foods and Beverages. Food Chem. 2020, 302, 125290. 10.1016/j.foodchem.2019.125290.

(35) Vargas, F.; Weldon, K. C.; Sikora, N.; Wang, M.; Zhang, Z.; Gentry, E. C.; Panitchpakdi, M. W.; Caraballo-Rodríguez, A. M.; Dorrestein, P. C.; Jarmusch, A. K. Protocol for Community-created Public MS/MS Reference Spectra within the Global Natural Products Social Molecular Networking Infrastructure. Rapid Commun. Mass Spectrom. 2020, 34 (10), e8725. 10.1002/rcm.8725.

(36) Greg Landrum, Paolo Tosco. Rdkit/Rdkit. https://github.com/rdkit/rdkit.

(37) Stincone, P.; Pakkir Shah, A. K.; Schmid, R.; Graves, L. G.; Lambidis, S. P.; Torres, R. R.; Xia, S.-N.; Minda, V.; Aron, A. T.; Wang, M.; Hughes, C. C.; Petras, D. Evaluation of Data-Dependent MS/MS Acquisition Parameters for Non-Targeted Metabolomics and Molecular Networking of Environmental Samples: Focus on the Q Exactive Platform. Anal. Chem. 2023, 95 (34), 12673–12682. 10.1021/acs.analchem.3c01202.

