## Supplemental information for "ModiFinder: Tandem Mass Spectral Alignment Enables Structural Modification Site Localization"

### **SI Note 1 - Alternative Evaluation Metrics for Localization Performance**

The first evaluation method that comes to mind is a binary classifier where it assigns 1 to the predicted array if the prediction likelihood assigned to the true modification site is the maximum prediction likelihood score within all atoms. This can be formulated as follows:

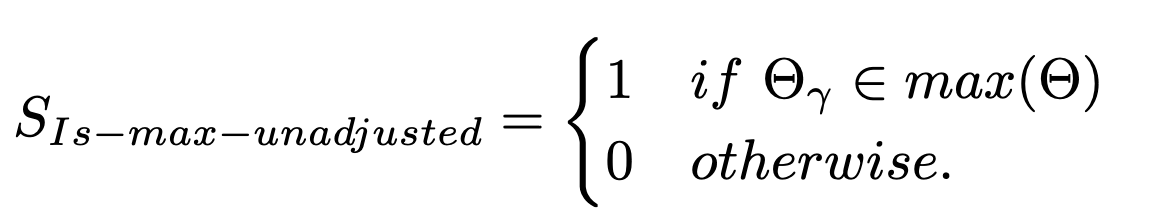

Where 𝚯 is the likelihood scores for the atoms, 𝜸 is the true modification site, and max(𝚯) is the set of all high-scoring atoms. This evaluation score falls short as an effective metric as it returns 1 in scenarios when every atom is assigned the same value, despite the array of predictions providing no valuable insights; highlighting its inadequacies for the ambiguity cover properties. Additionally, this method disregards the distribution of high-likelihood atoms in the structure, indicating it also falls short in the proximity cover properties as well. To adjust this metric for ambiguity, we propose the *Is-max* evaluation metric as follows:

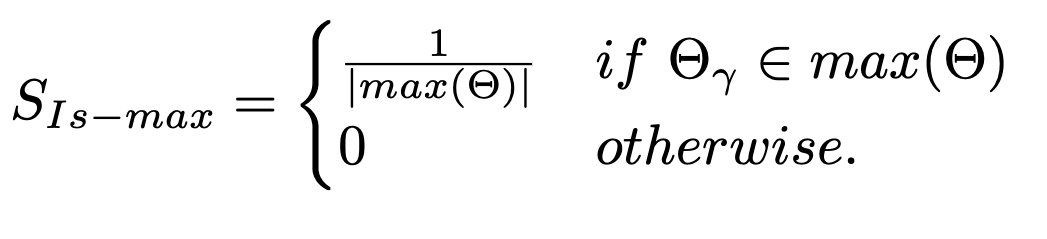

The next evaluation method is based on the shortest distance from the true modification site to an atom with the maximum prediction score. This can be formulated as:

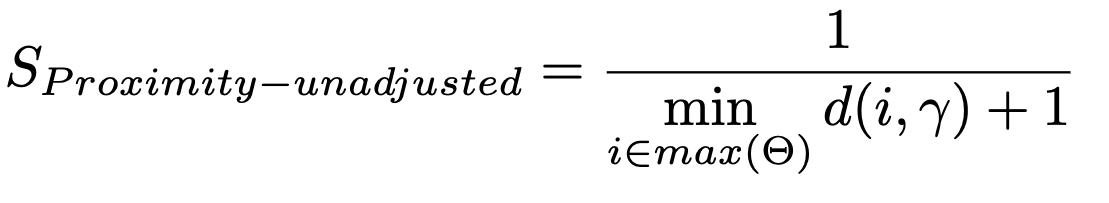

Where d(i, 𝜸) is the shortest distance from atom i to the true modification site on the graph. This method disregards the size of the atom completely and the absence of adjustment for the atom's overall size leads to a scoring bias, favoring smaller compounds with higher evaluation scores. A prediction indicating the furthest distance from the actual modification site in a small molecule may still be closer than a prediction targeting an atom within 10% of the true modification site, resulting in a higher score while providing less useful information. *Proximity method* adjusts this issue:

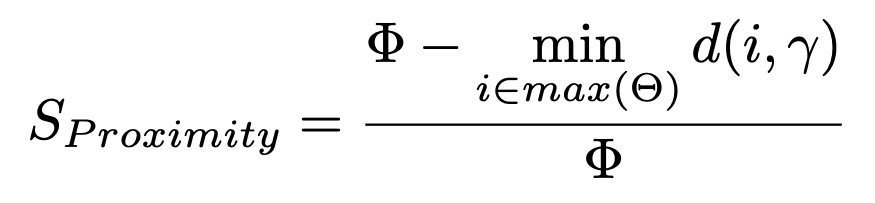

This method does not have a good ambiguity cover as the same example provided for *is-max-unadjusted* applies here as well. Our proposed evaluation method referred to as Average dist provides a balance between the ambiguity cover and proximity by taking both the scores and the relative distances into account.

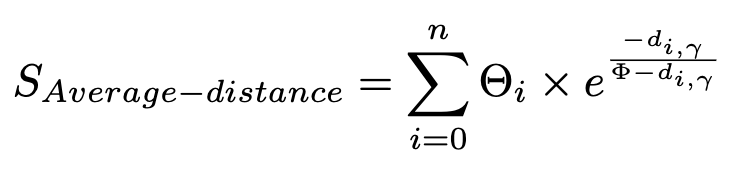

Along with these evaluation functions, a *Sorted-rank* evaluation is also reported as it has been used for similar applications before. The *Sorted-rank* is correlated to the relative location of the true modification site’s likelihood score in comparison to other atoms.

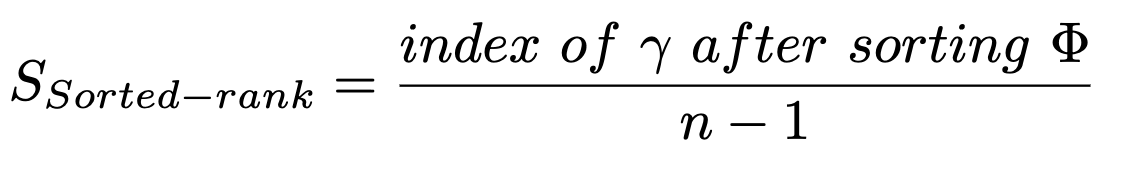

Where n is the number of atoms. If the true modification site has the highest score, it will be moved to the n-1th position after sort (index n) resulting in the evaluation score of 1. In contrast, if it has the lowest score, it will be moved to the first location after sorting (index 0) resulting in a score of 0. It is worth noting that the *Sorted-rank* also lacks proximity cover as it completely ignores the 2D structure graph.

#### **SI Note 2 - ModiFinder Imbalance Symmetry**

One of the scenarios that ModiFinder can fail is when there is a symmetry combined with an imbalance in the compound’s structure that leads to a bias. For example, assume the modification (M) in the toy example illustrated in **Figure SI-1 A** happens at the green puzzle piece A. In addition, assume there is a shifted peak corresponding to a fragment that has a green and a blue puzzle piece. In this case, Puzzle Piece B achieves a higher likelihood score as it appears in more potential structures (**Figure SI-1 B**).

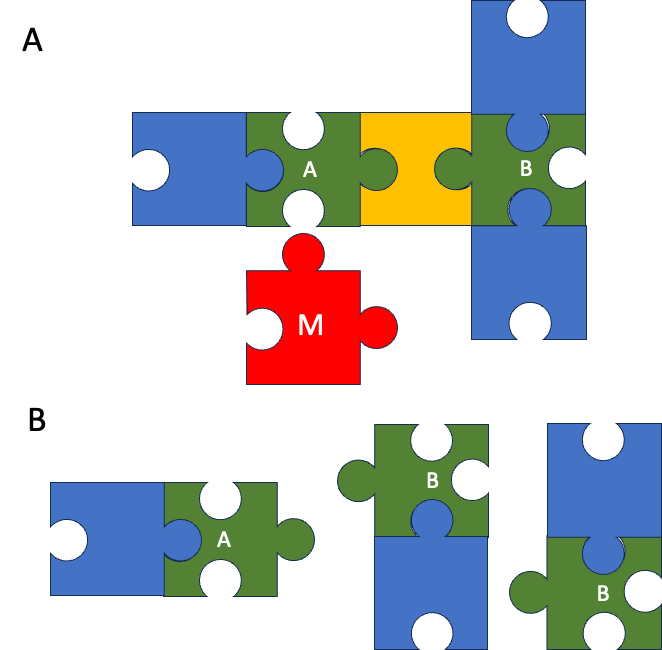

**Figure SI-1 -** **Illustration of** **imbalance symmetry on a toy example**. **A** This figure shows a base known compound and its modification (red piece M) and the modification site (Green piece A). **B** This shows all the substructures that have a green and a blue piece. Even though the modification happens at A and there is a potential substructure containing A, B achieves a higher likelihood score.

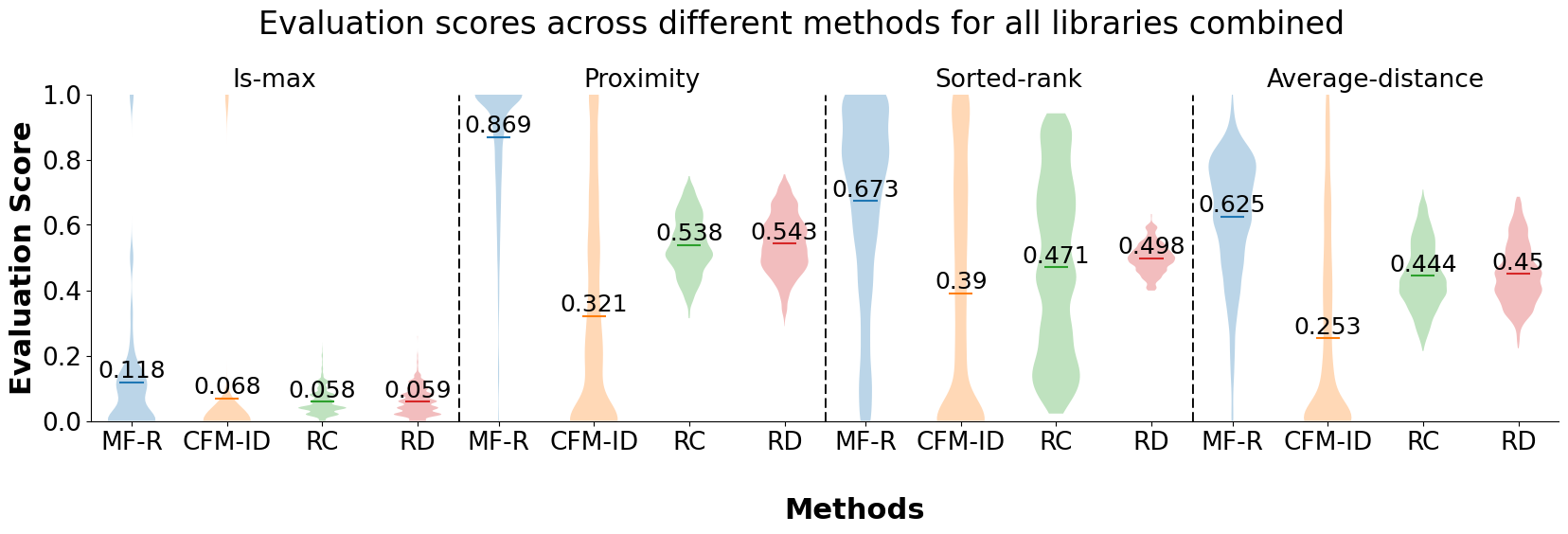

***Figure SI-2 - Comparative Analysis of Evaluation Scores:*** *Demonstrating the Consistent Superiority of ModiFinder Over Other Baselines Across Various Evaluation Methods.*

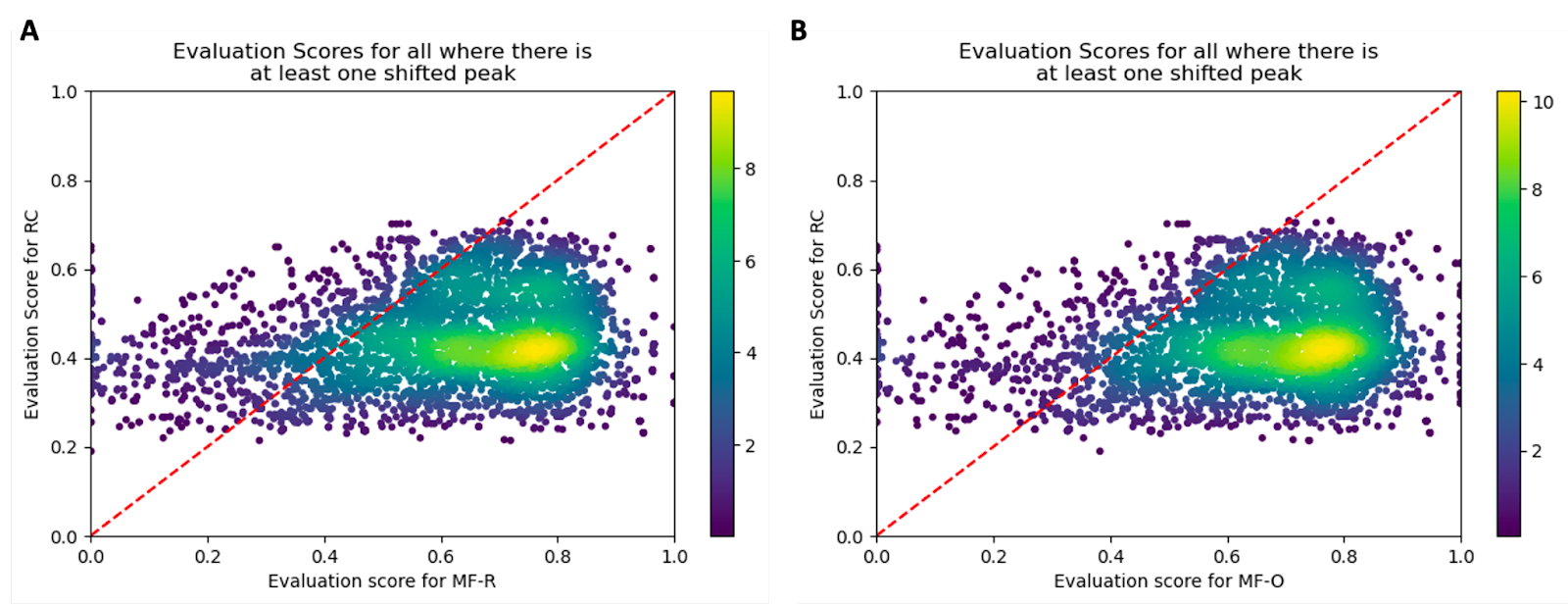

***Figure*** ***SI-3: Pairwise comparison of MF-R and MF-O with RC. A*** *Evaluation scores for all the pairs in the dataset using ModiFinder-Refined (MF-R) and Random Choice. The points below the dashed line represent instances where MF-R achieves a better resul than RC.* ***B:*** *Evaluation scores for all the pairs in the dataset using ModiFinder-Oracle (MF-O) and Random Choice calculated and plotted. The points below the dashed line represent instances where MF-R achieves a better result than RC.*

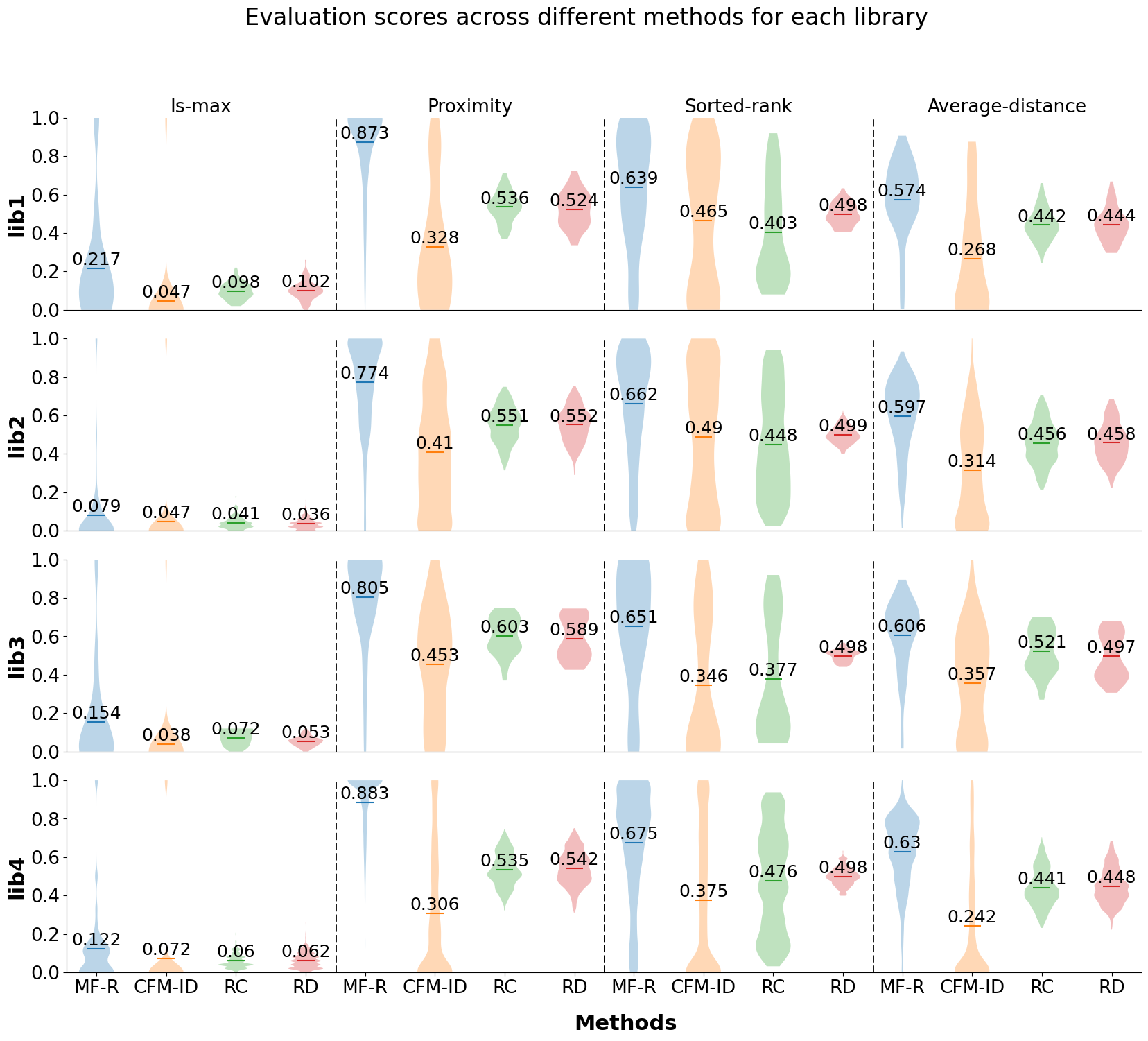

***Figure SI-4 Comparative Analysis of Evaluation Scores for each library:*** *Demonstrating the Consistent Superiority of ModiFinder Over Other Baselines Across Is-Max, proximity, Sorted-rank, and Average-distance evaluation method within each library.*

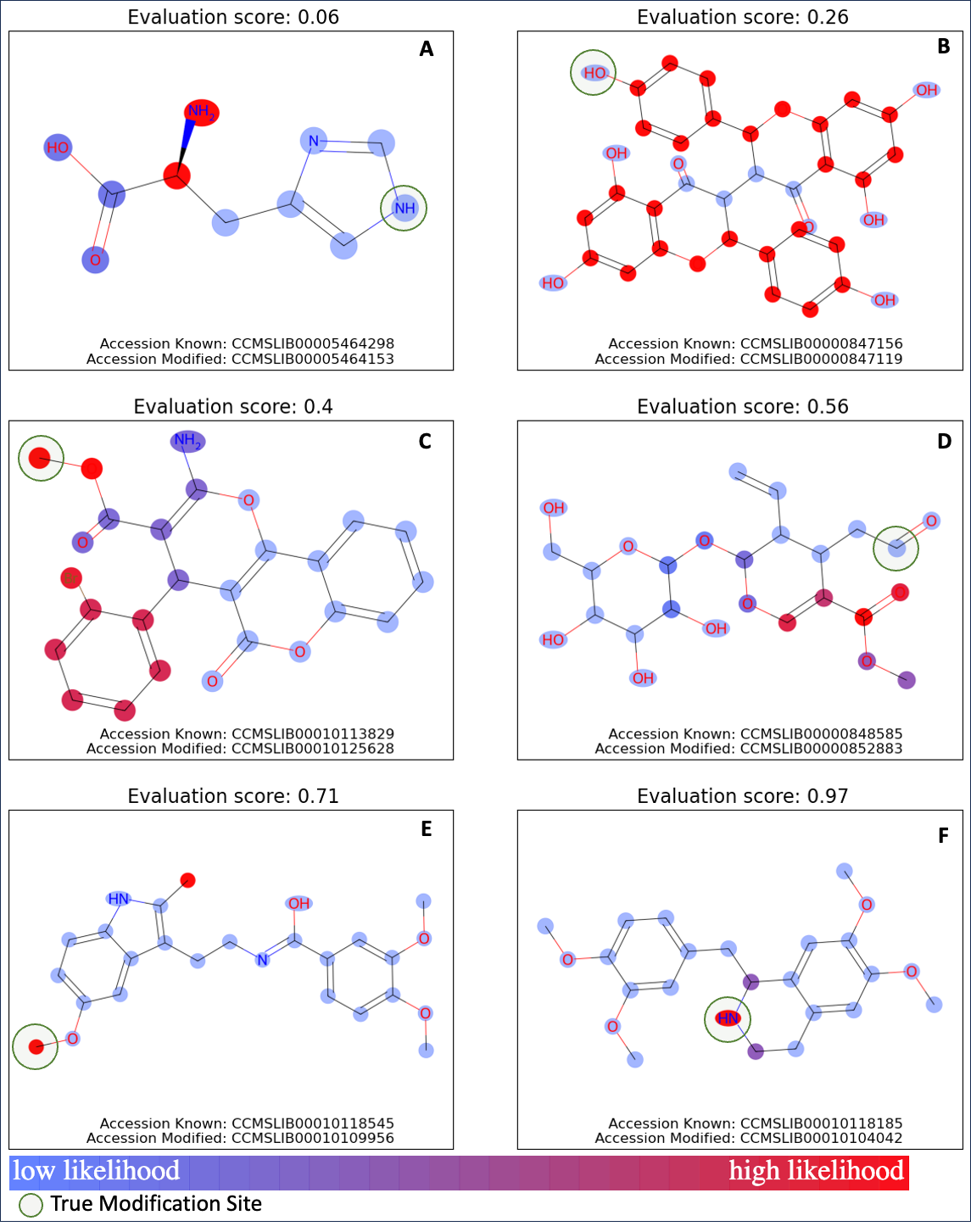

***Figure SI-5: The Evaluation score across different examples.*** *For each example, the evaluation score along is specified. The GNPS accession code of the known and the modified unknown compound is provided for each example. The true modification site is highlighted with the green circle.* ***A*** *The prediction has a bad proximity cover and receives the low score of 0.06.* ***B*** *This prediction has a bad ambiguity cover and receives the low score of 0.26.* ***C*** *This prediction is moderately ambiguous and therefore receives an evaluation score of 0.4.* ***D*** *This prediction cannot detect the true modification site and the prediction is somewhat distant leading the a reasonable score of 0.56 corresponding to its moderate proximity cover.* ***E*** *while modifinder can find the true modification site with low ambiguity, it is still penalized for the distance and ambiguity introduced by the second option leading to a score of 0.71.* ***F*** *The prediction has great proximity and low ambiguity leading to the high score of 0.97.*

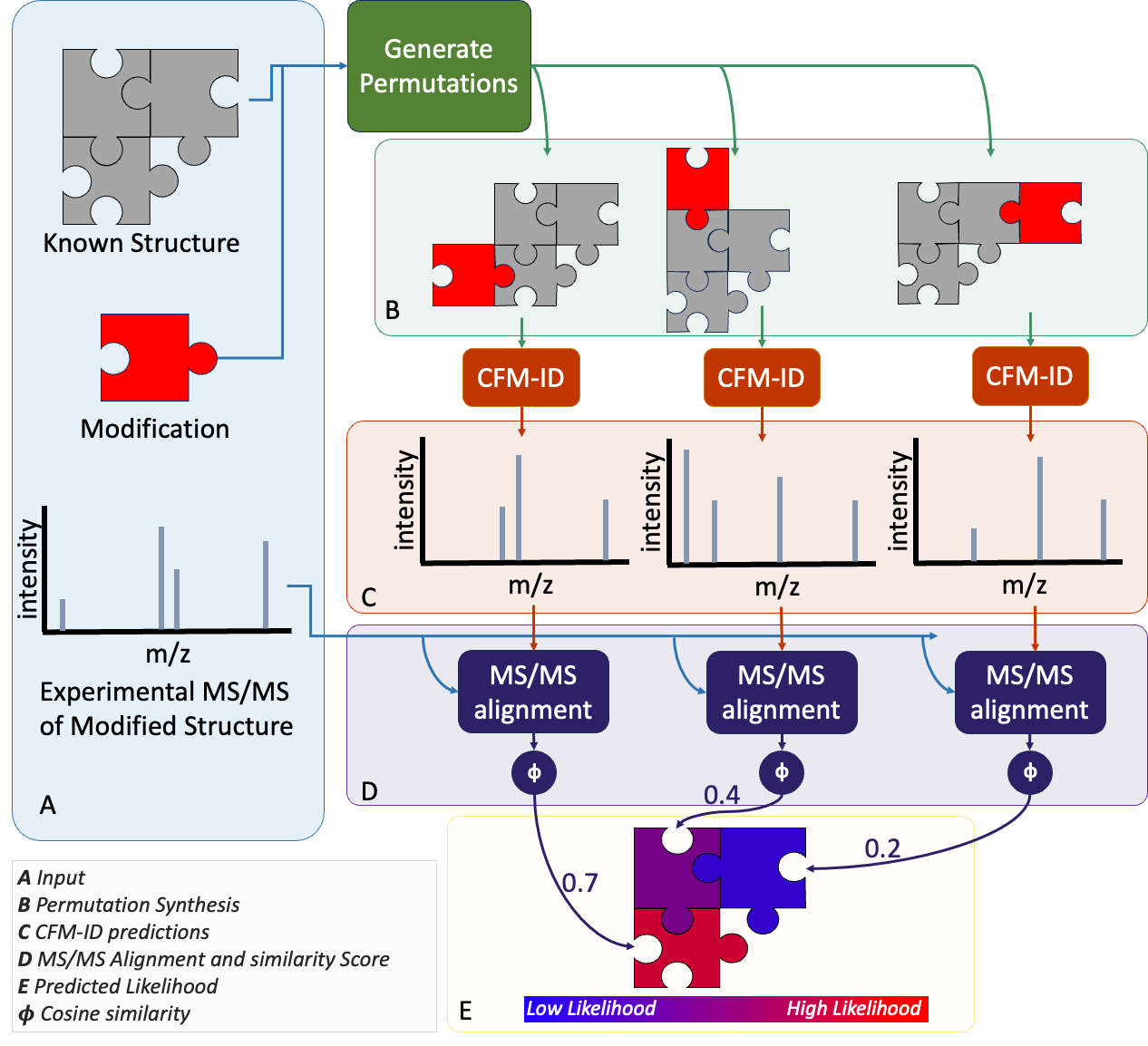

**Figure SI-6:** **Overview of the CFM-ID alternative approach**. **A.** the input section with two structures and the spectra of the bigger compound. **B.** permutation of the difference around the smaller compound to create possible analogs. **C.** prediction of the spectra using CFM-ID. **D.** Aligning the spectra of the larger molecule with each of the spectra predictions. **E.** assigning a likelihood score to each atom based on the cosine score of the alignment.

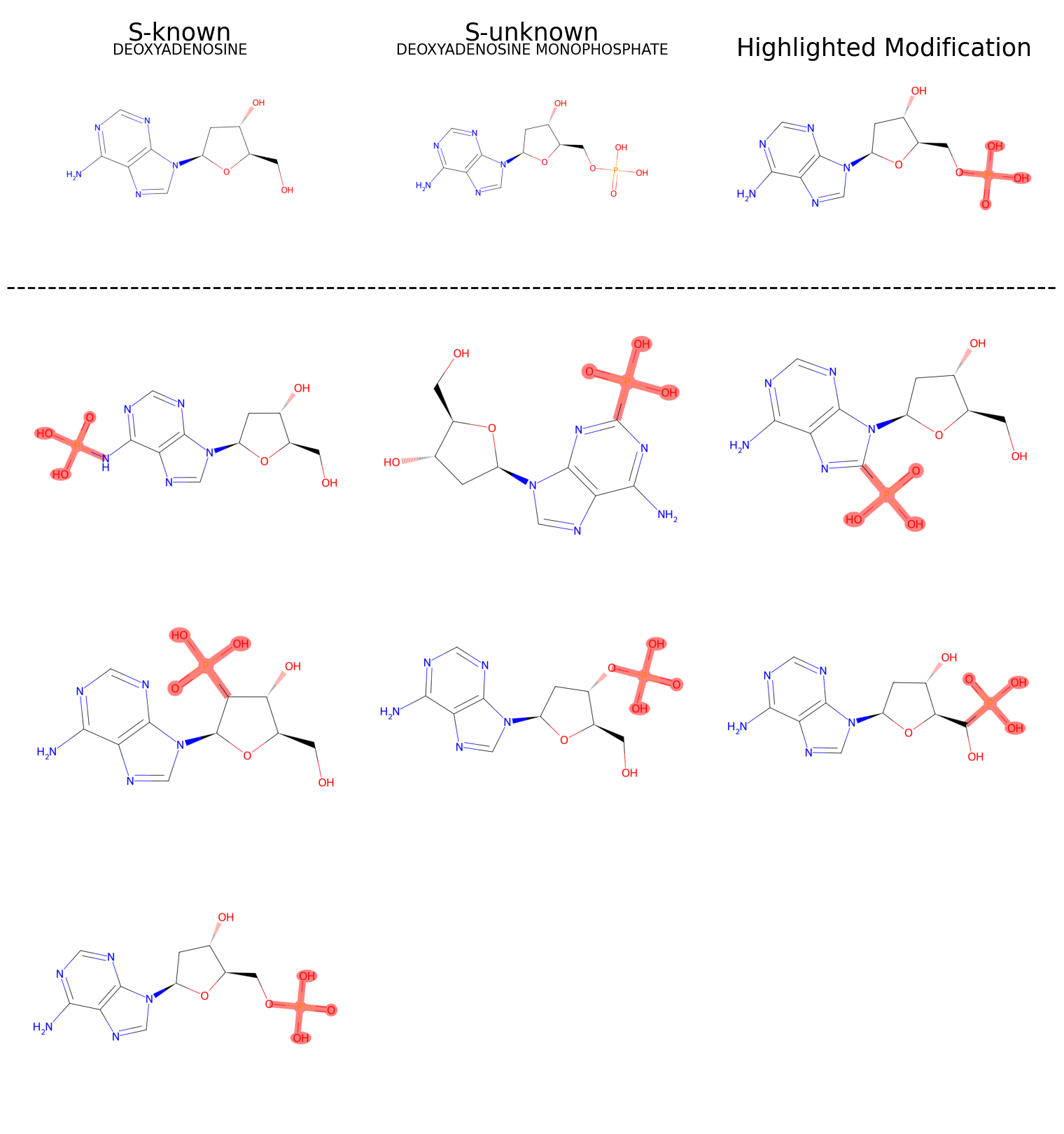

***Figure SI-7: Example of the permutation algorithm on Deoxyadenosine and Deoxyadenosine Monophosphate.*** *The upper section displays both molecules with the modification part highlighted. The lower section exhibits various molecules synthesized by permutating the modification on Deoxyadenosine.*

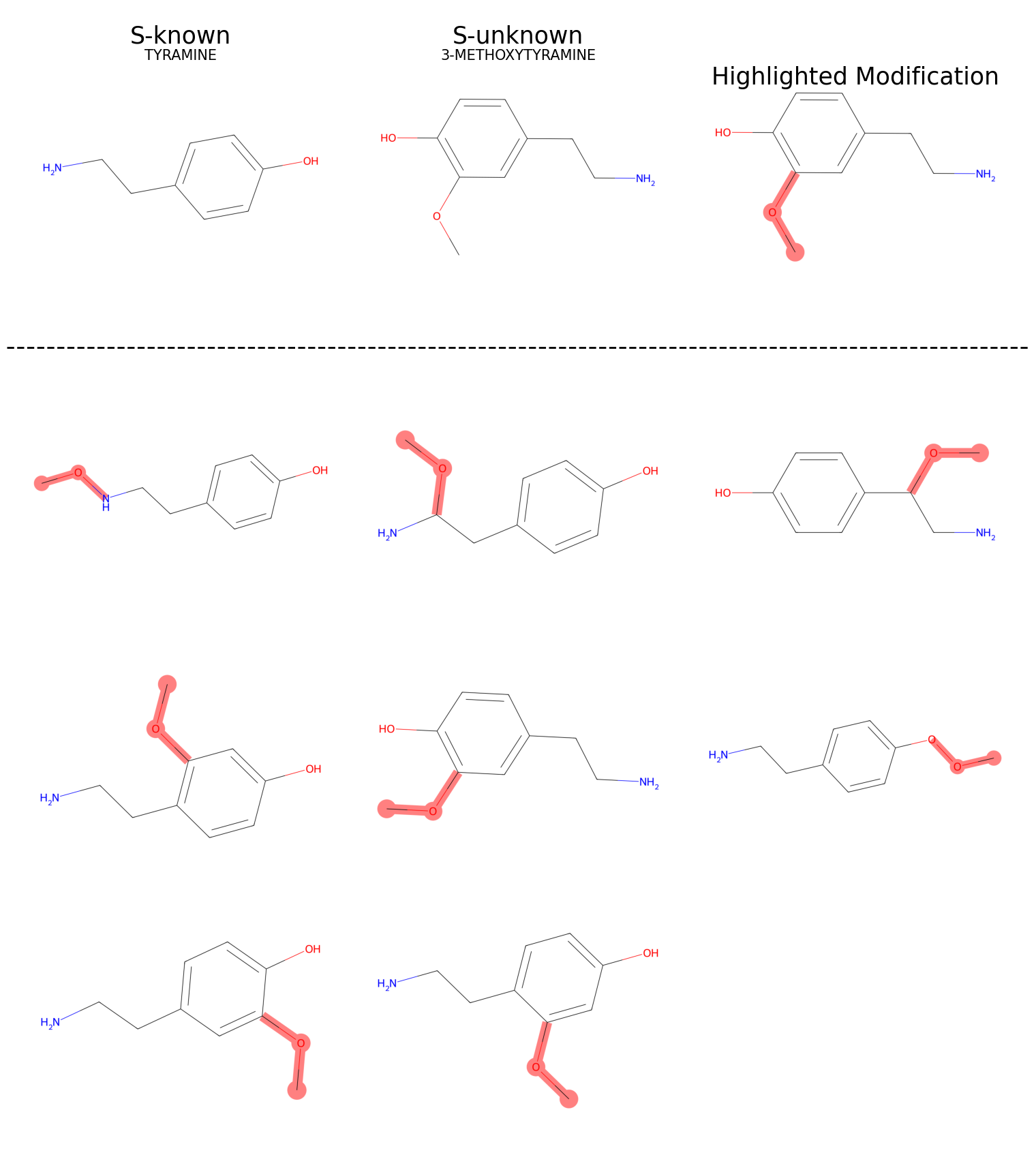

***Figure SI-8: Example of the permutation algorithm on Tyramine and 3-MethoxyTyramine.*** *The upper section displays both molecules with the modification part highlighted. The lower section exhibits various molecules synthesized by permutating the modification on Tyramine.*
